# PBK/TOPK mediates Ikaros, Aiolos and CTCF displacement from mitotic chromosomes and alters chromatin accessibility at selected C2H2-zinc finger protein binding sites

**DOI:** 10.1101/2024.04.23.590758

**Authors:** Andrew Dimond, Do Hyeon Gim, Elizabeth Ing-Simmons, Chad Whilding, Holger Kramer, Dounia Djeghloul, Alex Montoya, Bhavik Patel, Sherry Cheriyamkunnel, Karen Brown, Pavel Shliaha, Juan M. Vaquerizas, Matthias Merkenschlager, Amanda G. Fisher

## Abstract

PBK/TOPK is a mitotic kinase implicated in haematological and non-haematological cancers. Here we show that the key haemopoietic regulators Ikaros and Aiolos require PBK-mediated phosphorylation to dissociate from chromosomes in mitosis. Eviction of Ikaros is rapidly reversed by addition of the PBK-inhibitor OTS514, revealing dynamic regulation by kinase and phosphatase activities. To identify more PBK targets, we analysed loss of mitotic phosphorylation events in *Pbk^−/−^*preB cells and performed proteomic comparisons on isolated mitotic chromosomes. Among a large pool of C2H2-zinc finger targets, PBK is essential for evicting the CCCTC-binding protein CTCF and zinc finger proteins encoded by *Ikzf1*, *Ikzf3*, *Znf131* and *Zbtb11*. PBK-deficient cells were able to divide but showed altered chromatin accessibility and nucleosome positioning consistent with CTCF retention. Our studies reveal that PBK controls the dissociation of selected factors from condensing mitotic chromosomes and contributes to their compaction.

## Introduction

To improve our understanding of how cell identity is maintained when cells divide, there has been renewed interest in discriminating the factors that either remain associated with mitotic chromosomes or are displaced as chromosomes condense towards metaphase^1–13^. This discrimination is not entirely straightforward as fixation of mitotic samples has been shown to artificially strip some chromatin-associated factors from mitotic chromosomes^14^. In addition, although it is known that phosphorylation, mediated by mitotic kinases, can drive dissociation of proteins from mitotic chromosomes^15–23^ there is a paucity of knowledge regarding the extent to which factors are actively displaced, and which kinases, either alone or in combination, provoke the eviction of specific DNA-binding or chromatin-associated proteins. To close this knowledge gap, we have examined changes in the cell cycle distribution of key factors that regulate and determine cell commitment. Ikaros is a much studied lymphoid-restricted C2H2-zinc finger (ZF) transcription factor, encoded by the *Ikzf1* gene, that is critical for lymphocyte specification and differentiation, and has regulatory roles throughout the development of both B and T-cell lineages^24–30^. At the molecular level Ikaros forms dimers with itself or other close family members (such as Aiolos)^29,31–33^, and is postulated to either mediate silencing, or to activate transcription, according to context^34–37^. Ikaros proteins have also been shown to interact with the nucleosome remodelling and deacetylation complex (NuRD) in B and T-cell lineages^37–41^, and to bind at pericentric heterochromatin domains in lymphocytes during interphase^33,42–44^.

In a previous study^15^ Ikaros was shown to be inactivated in mitosis, potentially as part of a common mechanism shared with other proteins that contain C2H2-ZF DNA binding domains^15,16,45,46^. At the time this result was considered surprising because Ikaros is implicated in the heritable silencing of gene activity^43^ and could therefore be presumed to remain at sites of repression. Furthermore, prior studies by us and others have shown that many factors that contribute to durable chromatin-based epigenetic silencing remain chromosome-associated in mitosis^7,8,47–49^. This includes components of the DNA methylation machinery DNMT1,3A,3B; Polycomb Repressor Complexes 1 and 2; and the histone H3 lysine 9 methyltransferases SUV39H1/H2.

To verify that Ikaros does indeed dissociate from chromosomes during mitosis, and to better define the kinases responsible for this eviction, we generated tools that enabled real-time imaging of Ikaros distribution in living cells. We then used these to visualise Ikaros redistribution during the cell cycle and following treatment with a range of different kinase inhibitors. We discovered that PDZ-binding kinase (PBK, also known as T-lymphokine-activated killer-cell-originated protein kinase (TOPK)) is both necessary and sufficient to evict Ikaros from metaphase chromosomes, a finding confirmed by genetic knock-out (KO) of *Pbk*. Furthermore, treatment of Ikaros-depleted chromosomes with the PBK-inhibitor OTS514 provokes rapid re-association of Ikaros with condensed chromosomes, revealing dynamic regulation by kinase and phosphatase activities. To uncover other factors targeted for eviction by PBK/TOPK, we adopted a two-pronged approach. Firstly, we raised an antibody to a phosphorylated C2H2-ZF peptide linker and performed immunoprecipitation (IP) from mitotic lysates of B cell progenitors (preB cells) to screen for potential targets. Secondly, using a previously described approach^8,49^, we purified unfixed mitotic chromosomes from *Pbk^−/−^* or wildtype (WT, *Pbk^+/+^*) preB lymphocytes and compared their proteomes by quantitative liquid chromatography-tandem mass spectrometry (LC-MS/MS). This combined strategy, together with immunofluorescence (IF) comparisons, enabled us to identify and verify an important cohort of factors that are co-ordinately evicted from chromosomes by PBK. These include Ikaros, Aiolos, the DNA binding factors ZNF131, ZBTB11 and BCL11A, and the insulator protein CTCF. Chromatin accessibility data (using ATAC-seq) are consistent with greater retention of C2H2-ZF factors on *Pbk^−/−^*mitotic chromosomes, and we observe increased accessibility at multiple loci, especially at CTCF binding sites. *Pbk^−/−^* metaphase chromosomes are significantly less condensed that their normal counterparts, a result which suggests that PBK-mediated phosphorylation, and the subsequent displacement of C2H2-ZF factors, may be important to enable chromosomes to properly condense in mitosis.

## Results

### Ikaros is actively evicted from mitotic chromosomes by phosphorylation

Previous work has suggested that Ikaros binds to DNA in a cell-cycle-dependent manner^15,42,50^. Antibody-based staining of mouse preB cells confirmed that Ikaros localises to pericentric (heterochromatin) clusters in interphase, but is absent from mitotic chromosomes during metaphase, with association being re-established during mitotic exit (Figure 1a). This cell-cycle-dependent localisation of Ikaros was also seen in mouse T (VL3-3M2) and macrophage (J774A.1) cell lines (Supplementary Figures S1a and S1b) and is consistent with previous reports^42^. To independently verify that the mitotic eviction of Ikaros was not an artefact of sample fixation, we used CRISPR/Cas9 to engineer knock-in (KI) mouse preB cells that express Ikaros-mNeonGreen fusion proteins derived from the endogenous *Ikzf1* locus (Figure 1b). Two different gRNAs were used to generate four hetero- or homozygous KI clones expressing Ikaros-mNeonGreen fusion proteins (Figure 1b, middle/lower). Live-cell imaging of these clones revealed that Ikaros-mNeonGreen dissociates from condensing chromosomes during prophase and is no longer detected on chromosomes at metaphase (Figure 1c, Supplementary Figure S1c and Supplementary Videos S1-4). Ikaros-mNeonGreen showed a re-association with pericentric chromosomal regions during anaphase, and at telophase/cytokinesis. These results importantly validate our antibody-based observations and reveal that Ikaros re-association with chromosomes occurs rapidly towards the end of mitosis, prior to entry into G1 (Supplementary Videos S1-4).

**Figure 1.**
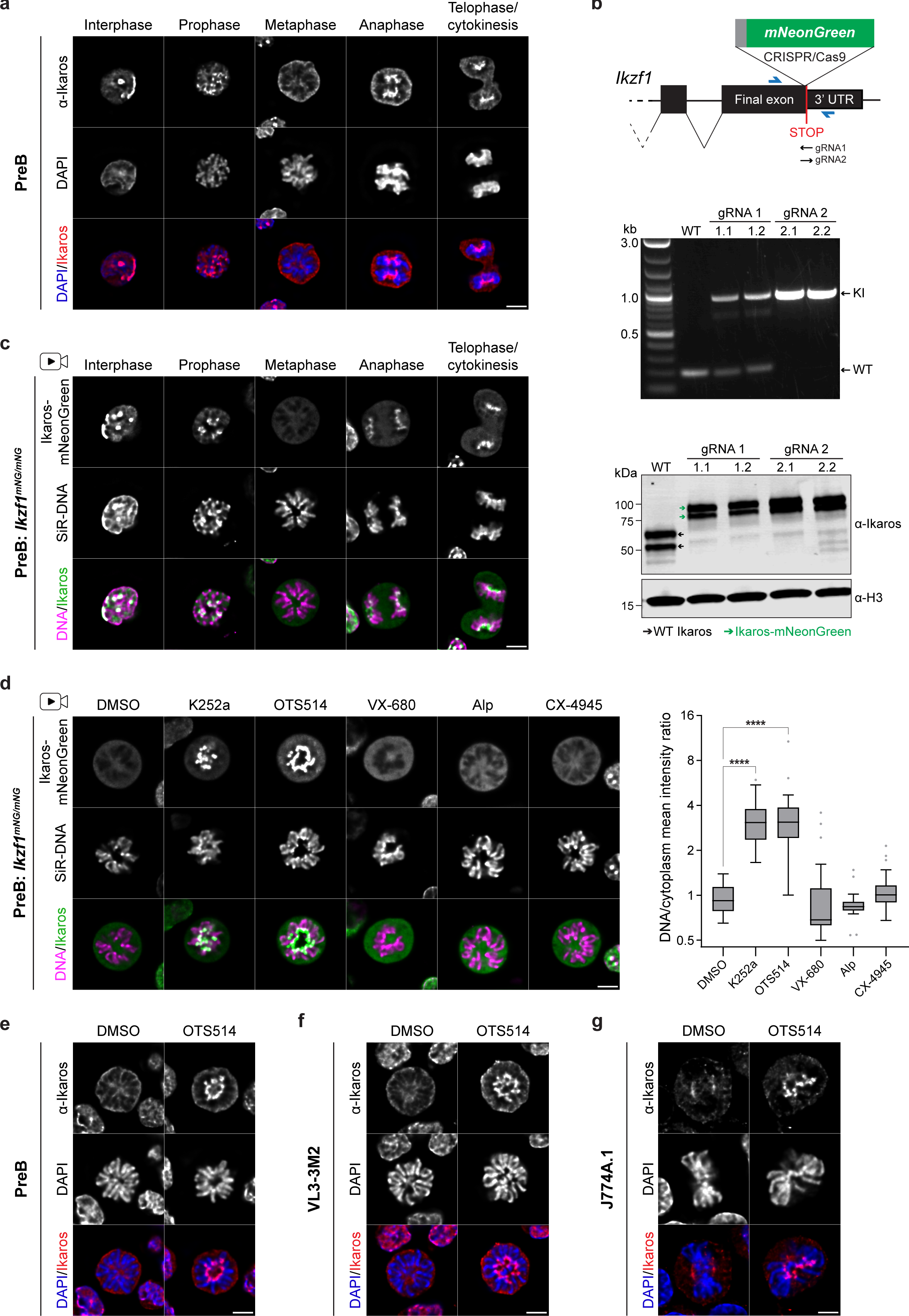
| Ikaros normally dissociates from metaphase chromosomes but can be retained following kinase inhibitor treatment. a. Immunofluorescence staining of Ikaros localisation in fixed mouse preB cells, showing representative images of interphase cells and progressive stages of mitosis. Scale bar=5 µm; at least 12 cells per mitotic stage (>50 for metaphase) were imaged across four independent staining experiments. b. Generation of Ikaros-mNeonGreen preB cells by CRISPR/Cas9 insertion of *mNeonGreen* immediately upstream of the endogenous *Ikzf1* STOP codon (illustrated top). Two different gRNAs were used in independent targeting experiments and knock-in (KI) was validated by PCR (middle, blue arrows in schematic illustrate primer positions), followed by sanger sequencing, in two selected clones from each targeting experiment. Clones 1.1 and 1.2 had *mNeonGreen* inserted into one allele and harboured frame-shift mutations in the apparent WT allele (*Ikzf1^mNG/–^*), whilst clones 2.1 and 2.2 were homozygous KI (*Ikzf1^mNG/mNG^*). Western blot (botom, anti-Ikaros (C-terminal) 1:5000) revealed all four clones expressed Ikaros-mNeonGreen fusion protein exclusively. c. Live-cell imaging of Ikaros-mNeonGreen in *Ikzf1^mNG/mNG^* homozygous KI mouse preB cells (clone 2.1), cultured with SiR-DNA, showing representative images of interphase cells and progressive stages of mitosis. Scale bar=5 µm; at least 11 cells per mitotic stage (>50 for metaphase) were imaged across four independent imaging experiments. d. Live-cell imaging of mitotic *Ikzf1^mNG/mNG^* mouse preB cells (clone 2.1) following 10 min treatment with DMSO or the indicated kinase inhibitors (K252a at 1 µM, all others at 10 µM; Alp=alsterpaullone). Images (left) are representative of mitotic cells from at least two independent treatment experiments (five replicates for K252a and OTS514). Cells were pre-cultured with SiR-DNA; scale bar=5 µm. Quantification (right) of mean chromosomal versus cytoplasmic mNeonGreen intensity is shown for one representative replicate of each treatment (quantification strategy illustrated in Supplementary Figure S1e; n≥18 cells per treatment; boxplots show median, interquartile range and Tukey whiskers, ploted on a log2 scale; ****padj<0.0001, Dunet’s multiple comparisons test). e. Immunofluorescence staining of Ikaros localisation in fixed mitotic preB cells following 10 min treatment with DMSO or 10 µM OTS514. Images are representative of three independent experiments, with bright centromeric staining observed in all OTS514 treated metaphase cells (63/63 for OTS514 versus 0/66 for DMSO). Scale bar=5 µm. f. Immunofluorescence staining of Ikaros localisation in fixed mitotic VL3-3M2 cells (mouse T cell line) following 10 min treatment with DMSO or 10 µM OTS514. Images are representative of three independent experiments, with 79% of OTS514 treated metaphase cells showing centromeric staining (n=total 80 cells; 34 bright, 29 mid, 17 low/none), compared to only 17% of DMSO treated cells showing any centromeric signal (n=total 75 cells; 0 bright, 13 mid, 62 low/none). Scale bar=5 µm. g. Immunofluorescence staining of Ikaros localisation in fixed mitotic J774A.1 cells (mouse macrophage cell line) following 10 min treatment with DMSO or 10 µM OTS514. Images are representative of two independent experiments, with bright centromeric staining observed in all OTS514 treated metaphase cells (42/42 for OTS514 versus 0/46 for DMSO). Scale bar=5 µm.

Previous literature suggests that phosphorylation can regulate the eviction of many DNA-binding factors in mitosis^15–23^. To better understand this, and to identify the mechanisms of Ikaros dissociation and re-association, actively dividing *Ikzf1^mNG/mNG^*preB cells were treated with a range of kinase inhibitors (Figure 1d and Supplementary Figure S1d; see Table 5 in Methods). By examining living cells during metaphase, we found that three inhibitors prevented Ikaros-mNeonGreen eviction from metaphase chromosomes, compared with DMSO-treated controls: K252a (a broad-spectrum kinase inhibitor; Figure 1d), and OTS514 and OTS964 (two related PBK inhibitors; Figure 1d and Supplementary Figure S1d). Cells treated with each of these inhibitors showed abundant Ikaros signal on metaphase chromosomes, particularly focused around centromeric domains. In contrast, treatment with other inhibitors, including those directed towards Aurora kinases (VX-680, Hesperadin), cyclin dependent kinases (Alsterpaullone (Alp)) or the known Ikaros kinase Casein kinase 2 (CX-4945)^51^, produced no appreciable change in Ikaros-mNeonGreen distribution as compared with controls (Figure 1d, left and Supplementary Figure S1d). Quantification of DNA/cytoplasmic signal ratios (Supplementary Figure S1e) confirmed that significant changes in Ikaros localisation were only seen following treatment with broad or PBK-specific inhibitors (Figure 1d, right). The capacity of OTS514 to block Ikaros eviction from metaphase chromosomes was verified using a second Ikaros-mNeonGreen KI clone (Supplementary Figure S1f), as well as by anti-Ikaros staining of normal preB cells expressing WT Ikaros (Figure 1e). Treatment of T cells and a macrophage cell line with OTS514 resulted in a similar retention of Ikaros on metaphase chromosomes (Figures 1f and 1g), consistent with a common mechanism of phosphorylation-induced mitotic eviction in different cell types.

### Ikaros distribution is dynamically regulated by competing PBK and phosphatase activities

Ikaros binding to chromosomes during the cell cycle appears highly dynamic, with rapid dissociation and re-association occurring during normal mitotic progression (Supplementary Videos S1-4). To determine whether the normal dissociation of Ikaros from chromosomes at metaphase could be reversed (rather than simply prevented) by PBK inhibition, individual mitotic cells in which Ikaros-mNeonGreen had already become dissociated were treated and monitored by live-cell imaging (Figure 2a). Immediately following addition of OTS514 inhibitor, increased signal was observed on at the centromeric regions of chromosomes, which rapidly increased over a matter of minutes (Figure 2a, quantified right). Given the speed of this re-association, we hypothesised that there might be an active mechanism to reverse Ikaros phosphorylation. In support of this, we showed that co-treatment with CalyculinA, a potent PP1/PP2A inhibitor, blocked OTS514-induced Ikaros re-association with mitotic chromosomes (Figure 2b). This result was also verified by antibody staining of Ikaros in WT preB cells (Supplementary Figure S2a). In contrast, Okadaic acid (OA), a PP2A phosphatase inhibitor with a lower potency for PP1, did not prevent PBK-inhibitor driven re-association of Ikaros (Figure 2b), suggesting PP1 may be responsible for Ikaros dephosphorylation.

**Figure 2.**
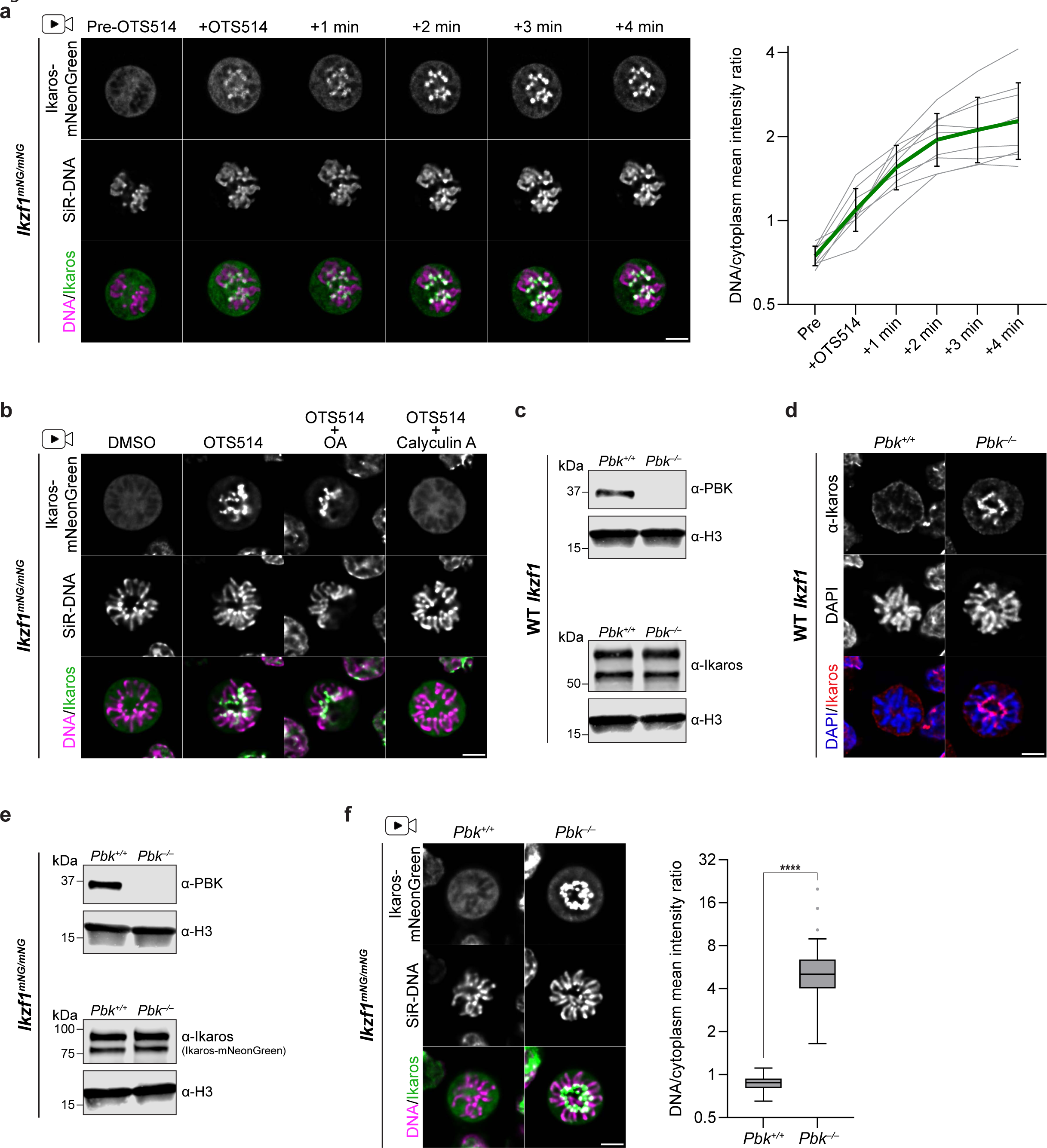
| Ikaros dissociation is regulated by competing PBK and phosphatase activities. a. Live-cell imaging of Ikaros-mNeonGreen re-localisation in individual mitotic *Ikzf1^mNG/mNG^* mouse preB cells (clone 2.1 from Figure 1b, pre-cultured with SiR-DNA) following addition of 10 µM OTS514. Representative live-cell imaging (left) of an individual mitotic cell before drug addition, immediately following treatment and during a further 4 minutes (scale bar=5 µm). Quantification (right) of mean chromosomal/cytoplasmic mNeonGreen signal for nine individual cells collected from six independent treatment experiments. Grey=individual cells; green=geometric mean; error bars=geometric standard deviation; ploted on a log2 scale. b. Live-cell imaging of mitotic *Ikzf1^mNG/mNG^* mouse preB cells (clone 2.1) following 10 min treatment with DMSO or with 10 µM OTS514 alone or in combination with the phosphatase inhibitors Calyculin A (100 nM) or Okadaic Acid (OA, 1 µM). Cells were pre-cultured with SiR-DNA; scale bar=5 µm. Images are representative of three independent replicates. Bright centromeric signal was observed in all OTS514 (25/25) and OTS514 + OA (28/28) treated metaphase cells, whereas 74% (20/27) of cells co-treated with Calyculin A showed no centromeric signal, with the majority of the rest (6/27) showing very weak signal. c. Western blot validation of CRISPR/Cas9-generated *Pbk^−/−^* mouse preB cells, confirming loss of PBK protein (upper, anti-PBK 1:1000) and similar levels of Ikaros protein (lower, anti-Ikaros(C-terminal) 1:1000) between *Pbk^+/+^* and *Pbk^−/−^* cells. d. Immunofluorescence staining of untagged Ikaros in fixed WT (*Pbk^+/+^*) or *Pbk^−/−^*mitotic preB cells. Images are representative of three independent staining experiments, with bright centromeric staining observed in all *Pbk^− /−^* metaphase cells (30/30 for *Pbk^−/−^* versus 1/54 with weak centromeric signal for *Pbk^+/+^*). Scale bar=5 µm. Results were further validated in a second PBK KO clone e. Western blot validation of *Pbk* KO in *Ikzf1^mNG/mNG^* mouse preB cells (clone 2.1), confirming loss of PBK protein (upper, anti-PBK 1:1000) and similar levels of Ikaros-mNeonGreen fusion protein (lower, anti-Ikaros(C-terminal) 1:1000) between *Pbk^+/+^* and *Pbk^−/−^* cells. f. Live-cell imaging of *Pbk^+/+^* or *Pbk^−/−^* mitotic mouse *Ikzf1^mNG/mNG^*preB cells cultured with SiR-DNA. Images (left) are representative of >70 mitotic cells from 3 independent imaging experiments. Scale bar=5 µm. Quantification (right) of mean chromosomal versus cytoplasmic mNeonGreen intensity is shown for one representative replicate (*Pbk^+/+^* n=32 cells, *Pbk^−/−^* n=34 cells; boxplots show median, interquartile range and Tukey whiskers, ploted on a log2 scale; ****p<0.0001, unpaired two-tailed t-test).

Although OTS514 and OTS964 are reported to be inhibitors of PBK, both can exhibit some off-target activity against other kinases^52,53^. Therefore, to confirm that PBK regulates the mitotic eviction of Ikaros from chromosomes, we undertook a genetic approach. PBK-KO (*Pbk^−/−^*) lines were generated from WT and *Ikzf1^mNG/mNG^* mouse preB cells using CRISPR/Cas9 (see Methods for details). PBK-null cells were capable of division and expressed Ikaros protein at levels comparable to the parental WT (Figure 2c). During interphase, *Pbk^−/−^* cells displayed a normal pattern of pericentric Ikaros staining (Supplementary Figure S2b). However, in mitosis we observed abnormal retention of Ikaros at metaphase chromosomes, with bright centromeric labelling in all mitotic *Pbk^−/−^* cells (Figure 2d). By targeting *Pbk* in preB KI cells expressing Ikaros-mNeonGreen (Figure 2e) we were also able to evaluate the impact of PBK elimination in live cells. *Ikzf1^mNG/mNG^ Pbk^+/+^* and *Pbk^−/−^* cells expressed comparable levels of Ikaros-mNeonGreen fusion protein (Figure 2e, lower), with signal localising to pericentric heterochromatin clusters in interphase, regardless of PBK status (Supplementary Figure S2c). However, as cells entered mitosis, Ikaros-mNeonGreen was seen to remain chromosome-associated in the absence of PBK (Figure 2f, quantified right). Interestingly, although treatment with CalyculinA was able to block the OTS514-indcued re-association of Ikaros (Figure 2b), it was insufficient to evict bound Ikaros in the absence of PBK (Supplementary Figure S2d). Together, these results show that Ikaros dissociation and re-association with mitotic chromosomes is dynamically and actively regulated through opposing kinase (PBK) and phosphatase (PP1) activities.

### PBK regulates mitotic chromosome compaction and protein composition

To determine whether PBK loss has wider impacts on mitotic chromosomes, beyond Ikaros retention, we used flow cytometry to purify native mitotic chromosomes^8,49^ from *Pbk^+/+^* and *Pbk^−/−^* mouse preB cells for downstream analysis (illustrated in Supplementary Figure S3a). We isolated two representative individual chromosomes, 3 and 19, from *Pbk^+/+^* and *Pbk^−/−^* cells (Supplementary Figure S3b and Figure 3a) and found that chromosomes (and centromeres) from PBK KO cells were significantly larger than equivalents isolated from WT cells (Figure 3b, quantified in Figure 3c). This increase in mitotic chromosome size in *Pbk^−/−^* cells was additionally verified using conventional metaphase spreads (Supplementary Figure S3c and Figure 3d) and shows that PBK activity is required for the normal compaction of mitotic chromosomes.

**Figure 3.**
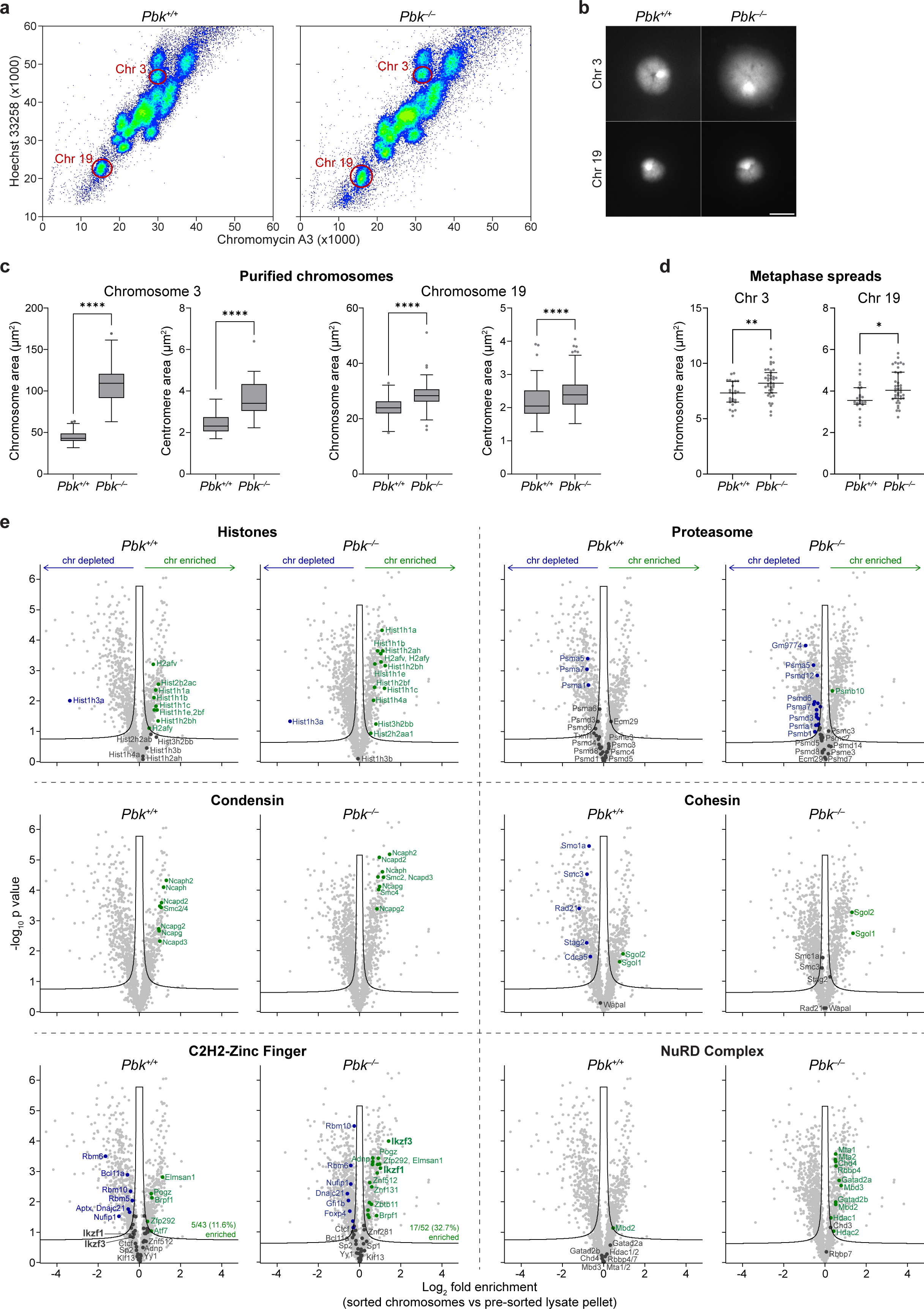
| Loss of PBK results in reduced mitotic chromosome compaction and increased retention of specific factors. a. Mitotic chromosomes from *Pbk^+/+^* and *Pbk^−/−^* mouse preB cells visualised by flow cytometry after staining with Hoechst 33258 and Chromomycin A3. Gates for purification of chromosomes 3 and 19 are shown; representative of three replicates. b. Flow-purified mitotic chromosomes 3 (upper) and 19 (lower) from *Pbk^+/+^* and *Pbk^−/−^* mouse preB cells, cytocentrifuged onto slides and stained with DAPI. Representative of chromosomes from three independent replicates; scale bar=5 μm. c. Size quantification of mitotic chromosomes 3 and 19, flow-purified from *Pbk^+/+^*and *Pbk^−/−^* mouse preB cells, measuring total (left) and DAPI-dense centromeric (right) areas. Quantification from one experiment, representative of three independent replicates (chromosome 3: n=117, 65 (total area) and n=96, 49 (centromeres); chromosome 19: n=133, 189 (total area) and n=113, 164 (centromeres); boxplots show median, interquartile range and Tukey whiskers; ****p<0.0001, unpaired two-tailed t-tests). d. Size quantification of chromosomes 3 and 19 in metaphase spreads from *Pbk^+/+^*and *Pbk^−/−^* mouse preB cells, identified by chromosome paints. Quantification from one experiment, representative of two replicates (n=26, 39 (chr 3) and n=24, 39 (chr 19); plots show median and interquartile range; **p=0.0017, *p0.05, unpaired two-tailed t-tests). e. Volcano plots highlighting specific factors enriched on or depleted from *Pbk^+/+^* and *Pbk^−/−^* mitotic chromosomes, compared to pre-sorted lysate pellets (green=enriched, blue=depleted, dark grey=not significantly enriched/depleted; modified two-tailed t-test with permutation-based false discovery rate (FDR)<0.05, S0=0.1; n=4 chromosome samples and n=3 lysate pellet samples; selected annotations are shown for proteasome and C2H2-zinc fingers; volcano plots highlighting all enriched/depleted factors are shown in Supplementary Figure S3f).

PBK has previously been implicated in regulating multiple targets in mitosis, particularly other C2H2-ZF transcription factors^46^. To investigate whether loss of PBK affects retention of other C2H2-ZF factors by mitotic chromosomes, or more broadly other chromatin components, we isolated total mitotic chromosomes from *Pbk^+/+^* and *Pbk^−/−^* mouse preB cells (Supplementary Figure S3d), and subjected these to proteomics analysis, as previously described^8,49^ (and detailed in Methods). To determine the factors enriched on mitotic chromosomes in each condition, we performed LC-MS/MS analysis of sorted chromosomes, compared to pre-sorted total mitotic lysate pellets (as illustrated in Supplementary Figure S3a). Replicates were similar for both *Pbk^+/+^* and *Pbk^−/−^* conditions, whilst sorted and unsorted samples clustered separately (Supplementary Figure S3e). Similar numbers of proteins were detected in WT and KO conditions, with a comparable proportion showing significant enrichment (17.1% versus 17.8%) or depletion (25.2% versus 29.9%) from mitotic chromosomes (Supplementary Figure S3f).

As expected, both *Pbk^+/+^* and *Pbk^−/−^*mitotic chromosomes showed an enrichment for histone proteins (Figure 3e, top left) and condensin complex proteins (Figure 3e, middle left), while components of the proteasome were not significantly enriched (Figure 3e, top right). Amongst cohesin complex components, only Shugoshin proteins (Sgo1/2) were significantly enriched on *Pbk^+/+^* and *Pbk^−/−^*chromosomes (Figure 3e, middle right), consistent with the general eviction of cohesin during prophase (although depletion was less pronounced in *Pbk^−/−^*samples), and the selective protection of centromeric cohesin by Shugoshin until anaphase^54^. Inspection of C2H2-ZF factors revealed that most, including Ikaros (Ikzf1) and Aiolos (Ikzf3), were either depleted or showed no significant enrichment on mitotic chromosome samples derived from WT (*Pbk^+/+^*) cells (Figure 3e, lower left). In *Pbk^−/−^* samples, we detected more C2H2-ZF factors overall (52 versus 43) and the proportion showing co-enrichment with chromosomes nearly tripled. Ikaros and Aiolos were among the factors enriched exclusively on *Pbk^−/−^* chromosomes, whilst other factors such as BCL11A no longer showed significant depletion in the absence of PBK. Despite an increase in mitotic chromosome retention of multiple C2H2-ZF factors in *Pbk^−/−^*samples, approximately two-thirds remained as not significantly enriched. These results suggest that PBK activity is required for the dissociation of a subset of C2H2-ZF proteins from mitotic chromosomes, whilst others can be evicted by alternative mechanisms.

To further examine changes in the association of factors with *Pbk^−/−^*mitotic chromosomes, we performed gene ontology (GO) analysis of depleted or enriched factors (Supplementary Figure S3g). Among the factors defined as depleted from *Pbk^+/+^* and *Pbk^−/−^* chromosomes, there was a broad similarity in overrepresented terms (Supplementary Figure S3g, lower), with spliceosomal components showing mitotic depletion in both conditions. However, among the proteins defined as enriched, we observed differences in overrepresented terms between *Pbk^+/+^* and *Pbk^−/−^*samples (Supplementary Figure S3g, upper). This included a modest reduction in outer-kinetochore proteins, a small increase in heterochromatin-associated proteins, and an apparent enrichment of nucleosome remodelling complexes, including ISWI-, SWI/SNF superfamily-, INO80- and CHD-type complexes, and exemplified by the NuRD complex (Figure 3e, lower right). Taken together, these results indicate that PBK activity is required for the mitotic eviction of a subset of C2H2-ZF proteins, may affect the normal dissociation of nucleosome remodelling complexes, and is important for normal chromosome compaction.

### PBK directly phosphorylates multiple C2H2-ZF proteins and regulates mitotic dissociation of CTCF

To further investigate potential targets of PBK, we generated a specific anti-phospho-linker antibody. Ikaros has previously been shown to be phosphorylated at specific sites within the highly conserved C2H2-ZF linkers^15,50^, and we therefore used a phosphorylated Ikaros136-143 linker peptide to generate rabbit antisera (as illustrated in Figure 4a, see Table 4 in Methods for details). This antibody detected an array of proteins in mitotic lysates from WT preB cells which were only weakly detectable in asynchronous lysates, and which were not detected in *Pbk^−/−^*samples (Figure 4b). Enhanced phosphorylation of histone H3 at serine 10 (H3S10p) was detected in mitotic versus asynchronous samples as expected, but levels were comparable in *Pbk^−/−^* and *Pbk^+/+^* samples, despite previous suggestions that PBK may contribute to H3S10 phosphorylation^55^. To identify proteins that were recognised by the anti-phospho-linker antibody, we performed quantitative proteomic comparisons of *Pbk^+/+^* and *Pbk^−/−^* mitotic lysates following immunoprecipitation (IP). Although protein abundances in total mitotic lysates appeared similar between conditions (Supplementary Figure S4a), we detected a wide range of factors in WT samples following IP with the anti-phospho-linker antibody, which were absent or significantly reduced following IP from *Pbk^−/−^* samples (Figure 4c, high/low values are shown by a blue/yellow scale). Overall, 111 hits were detected solely, or significantly more abundantly, in precipitates derived from *Pbk^+/+^*lysates, as compared to *Pbk^−/−^* lysates. Whilst unlikely to be an exhaustive list of substrates, this provides a catalogue of putative mitotic phosphorylation targets of PBK. The majority of these factors (72%) correspond to C2H2-ZF proteins (highlighted in red, Figure 4c), including SP1, YY1, CTCF, Ikaros and Aiolos. The differential detection of phosphorylated proteins between *Pbk^−/−^* and *Pbk^+/+^* samples was not caused by an underlying change in cell-cycle distribution or a reduced efficiency of mitotic arrest in mutant cells (Supplementary Figure S4b). Furthermore, since acute treatment of mitotically arrested WT preB cells with OTS514 resulted in a loss of antibody-reactive proteins (Supplementary Figure S4c), these results are consistent with a requirement for PBK to mediate and maintain phosphorylation of a cohort of proteins in mitosis.

**Figure 4.**
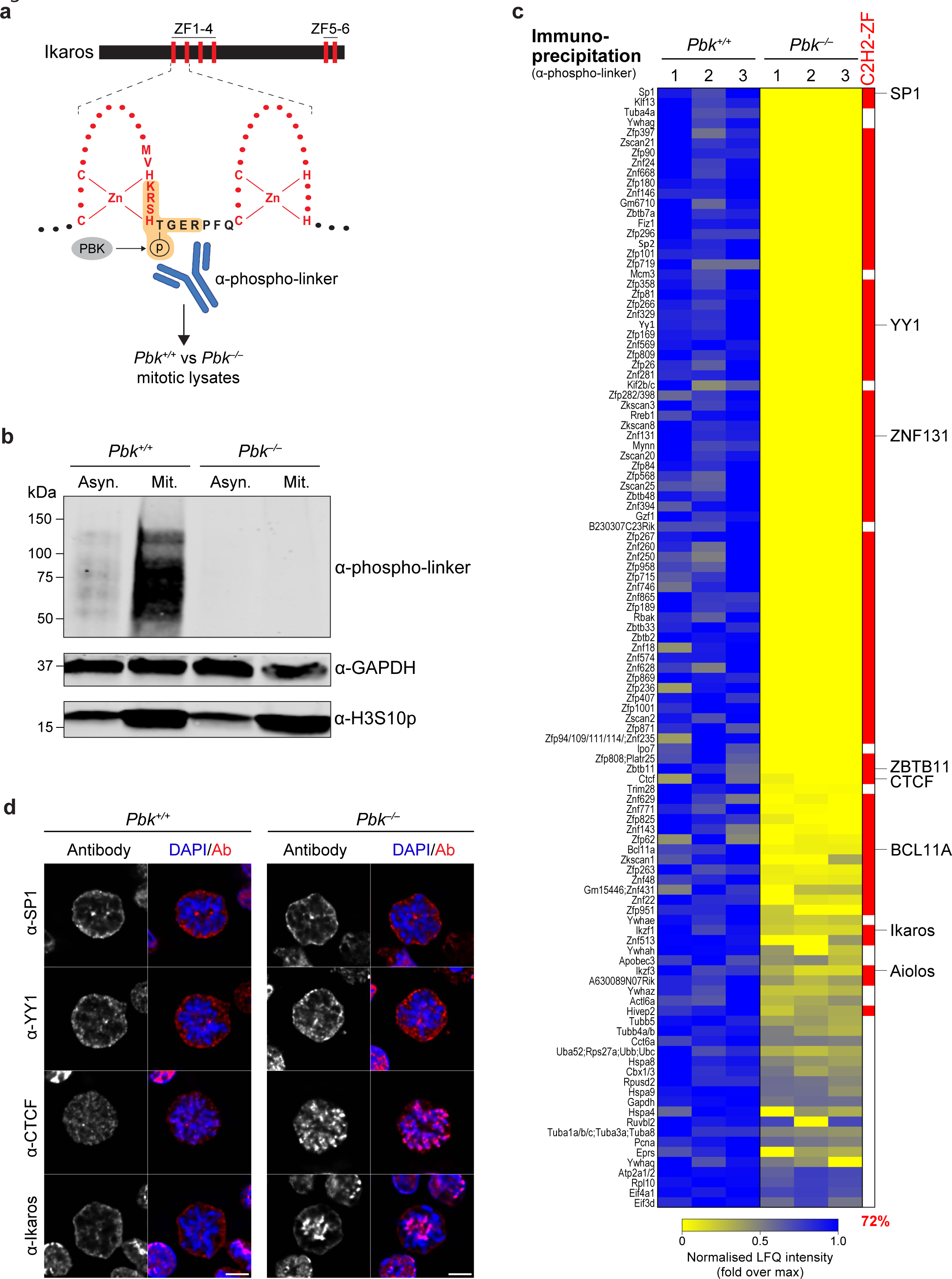
| PBK phosphorylates and regulates multiple C2H2-ZF proteins in mitosis. a. Schematic of Ikaros protein illustrating positions of zinc fingers (ZFs) 1-4, involved in DNA binding, and ZFs 5-6, involved in dimerization. ZFs 1 and 2 (red) and the conserved linker sequence (black) are shown in more detail, along with the proposed PBK mitotic phosphorylation site^15^. An anti-phospho-linker antibody was generated against a phosphorylated peptide corresponding to the shaded sequence and used to probe mitotic lysates from *Pbk^+/+^* or *Pbk^−/−^* mouse preB cells. b. Western blot detection of proteins recognised by anti-phospho-linker antibody in asynchronous (Asyn.) and mitotically arrested (Mit.) *Pbk^+/+^* and *Pbk^−/−^* mouse preB cell lysates. GAPDH was used as a loading control and H3S10p was included as a mitotic marker. Representative of three independent experiments, with results further validated in a second PBK KO clone. c. Heatmap representation of LC-MS/MS analysis of proteins immunoprecipitated with anti-phospho-linker antibody from *Pbk^+/+^* mitotic lysates, compared to *Pbk^−/−^* control mitotic lysates (n=3+3). Normalised LFQ intensity values, for three independent replicates, are shown for factors (111) which were detected after IP from *Pbk^+/+^*mitotic lysates, but which were undetected or at significantly lower levels after IP from *Pbk^−/−^* lysates (modified two-tailed t-test with permutation-based FDR<0.05 and S0=0.1; see Methods for full filtering criteria). Proteins containing a C2H2-ZF domain are highlighted in red. d. Immunofluorescence staining of the indicated factors in fixed WT (*Pbk^+/+^*) or *Pbk^−/−^* mitotic preB cells. Images are representative of >16 mitotic cells from across two (YY1, CTCF) or three (SP1, Ikaros) independent staining experiments. Staining paterns were further validated in a second PBK KO clone (SP1, CTCF, Ikaros) and with an alternative antibody (SP1). Ikaros staining is derived from the experiment shown in Figure 2d and a further representative image is included here for comparison. Scale bar=5 µm.

We next compared PBK targets identified by anti-phospho-linker IP with those identified by proteomic analysis of flow-sorted mitotic chromosomes. While there was only a modest overlap in these datasets, more C2H2-ZF PBK targets were detected overall in *Pbk^−/−^* chromosome samples than in *Pbk^+/+^* samples (Supplementary Figure S4d), consistent with a greater retention of these factors in the absence of PBK activity. It was noticeable however that despite a loss of phosphorylation, there was not evidence of a wholescale redistribution of C2H2-ZF phosphorylation targets in the absence of PBK, and most of these factors were not significantly enriched on *Pbk^−/−^* chromosomes (Supplementary Figure S4d). Instead, a small subset of PBK phosphorylation targets, namely Ikaros, Aiolos, ZNF131 and ZBTB11, showed a significant enrichment exclusively on *Pbk^−/−^* chromosomes, while BCL11A was no longer depleted. Immunofluorescence analysis of selected C2H2-ZF factors in *Pbk^+/+^* and *Pbk^−/−^* cells confirmed that whereas Ikaros requires PBK for mitotic dissociation, release of SP1 and YY1 is independent of PBK phosphorylation (Figure 4d). In contrast, CTCF showed increased retention on the arms of mitotic *Pbk^−/−^* chromosomes in metaphase and anaphase (Figure 4d and Supplementary Figure S4e), whereas in WT *Pbk^+/+^* preB cells CTCF largely dissociates from chromosomes by metaphase and re-associates with chromosomes during telophase/cytokinesis (Supplementary Figure S4e). Interestingly, this factor was not previously identified as being significantly enriched by proteomics, which could reflect technical limitations associated with this approach, such as dynamic flux. Collectively our data indicate that PBK-mediated phosphorylation is necessary and sufficient for the mitotic eviction of a subset of C2H2-ZF transcription factors that includes Ikaros, Aiolos, ZNF131, ZBTB11, BCL11A and CTCF.

### Loss of PBK alters mitotic chromatin accessibility at C2H2-ZF protein binding sites

We have shown that mitotic chromosomes isolated from *Pbk^−/−^*cells preB cells were less compact (Figures 3b-d) and more enriched for certain C2H2-ZF binding proteins (Figure 3e, lower left; and Figure 4d) than WT equivalents. To understand the impacts of PBK-mediated phosphorylation on chromatin accessibility, we performed ATAC-seq on isolated native mitotic chromosomes from *Pbk^+/+^* and *Pbk^−/−^* cells, that were purified by flow cytometry to ensure high mitotic purity irrespective of synchronisation efficiency (illustrated in Figure 5a). Fragment size distributions, corresponding to both nucleosome-free and nucleosomal fragments, were very similar between conditions (Supplementary Figure S5a). Overall, chromatin accessibility was also broadly similar, although *Pbk^+/+^*and *Pbk^−/−^* samples clearly segregated by principal component analysis (Supplementary Figure S5b). Differential accessibility analysis revealed ∼15,000 differentially accessible peaks (Figure 5b), of which most (11,479 versus 3,811) showed increased accessibility in the absence of PBK. Regions of increased accessibility are strongly biased towards transcription start sites (TSSs), compared to the distribution of peaks which are unchanged or lose accessibility (Figure 5c). By examining individual genomic loci, we confirmed that accessibility was largely unchanged in the absence of PBK at most regions, including at *Ikzf1* (Supplementary Figure S5c, upper) and, perhaps surprisingly, at known Ikaros target genes such as *Igll* and *Vpreb1*^34–36,41^ (Supplementary Figure S5c, lower). However, specific differences were evident at some loci (Figure 5d), including gains of accessibility at previously inaccessible regions and more modest alterations at peaks already present in WT samples. Many increased peaks were located at TSSs (Figures 5c-d) which were enriched for genes involved in mRNA processing, translation, and protein degradation (Supplementary Figure S5d, upper); whilst genes with decreased TSS accessibility showed minimal GO term enrichment (Supplementary Figure S5d, lower). We also observed that many sites gaining ATAC-seq signal appeared to overlap with known CTCF binding sites (Figure 5d). To further investigate factors which might be responsible for this differential accessibility, we performed motif enrichment analysis at differentially accessible sites. Peaks with increased accessibility in *Pbk^−/−^* samples were highly enriched for CTCF and other C2H2-ZF motifs (Figure 5e). In contrast, peaks with decreased accessibility were generally not enriched for C2H2-ZF motifs (Supplementary Figure S5e); although interestingly, the only C2H2-ZF representative amongst the top ten enriched motifs at these sites corresponded to the repressor ZBTB11, which was found to require PBK phosphorylation for mitotic eviction (Figure 4c and Supplementary Figure S4d).

**Figure 5.**
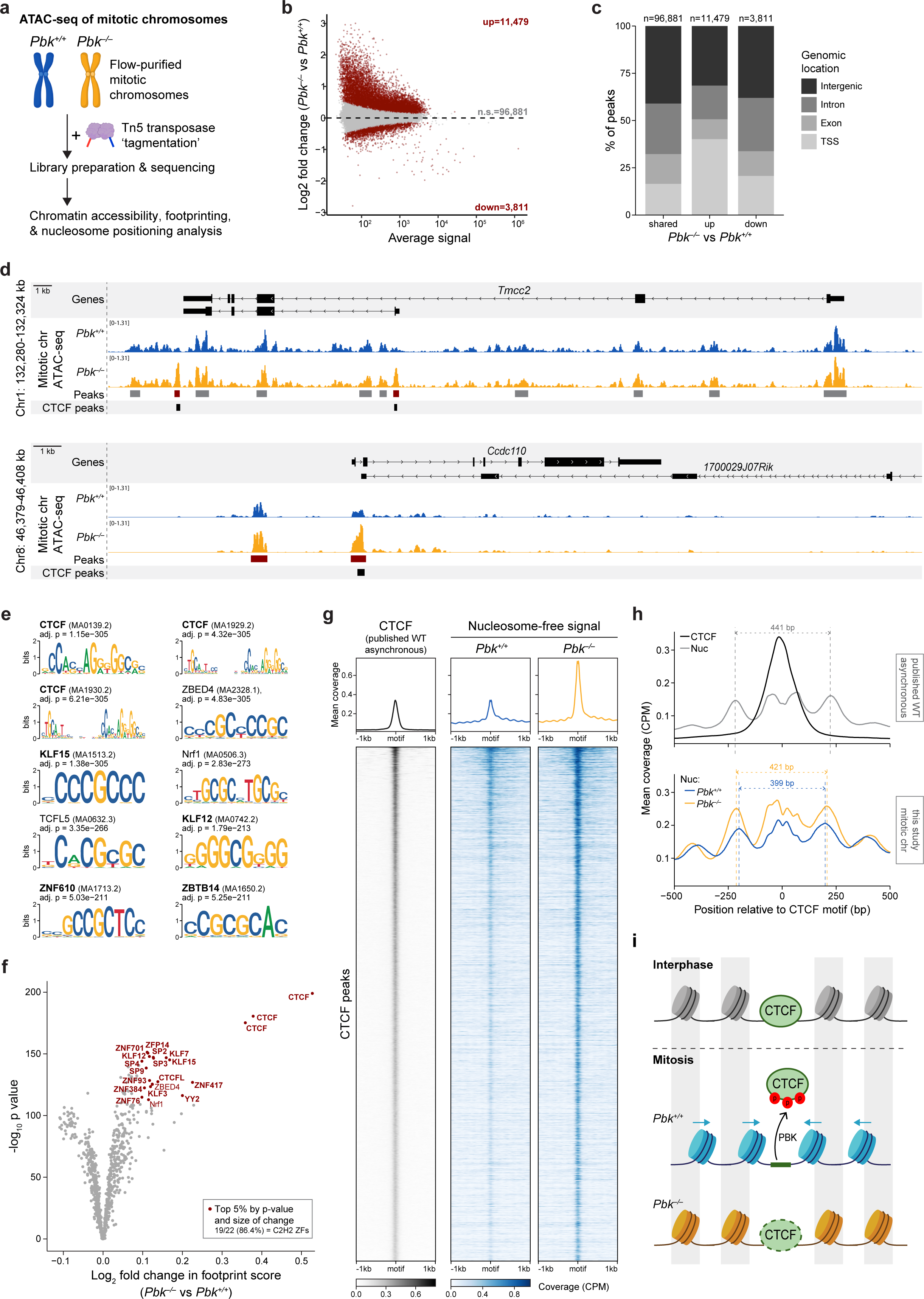
| Mitotic chromosomes from *Pbk^−/−^* cells have higher chromatin accessibility and evidence of increased CTCF retention. a. Illustration of experimental approach to assess flow-purified mitotic chromosomes from *Pbk^+/+^* and *Pbk^−/−^* mouse preB cells by ATAC-seq. b. MA plot showing differential accessibility analysis of *Pbk^−/−^* versus *Pbk^+/+^* mitotic chromosomes. Analysis was performed in DESeq2 using ATAC-seq counts at consensus MACS2 peaks, with significantly altered peaks highlighted in red (n=4+4; padj<0.1). c. Genomic distribution of ATAC-seq peaks which were unchanged or showed significant changes in accessibility between *Pbk^+/+^* and *Pbk^−/−^* mitotic chromosomes. d. Representative loci showing nucleosome-free (≤100bp) ATAC-seq signal from *Pbk^+/+^*(blue) and *Pbk^−/−^* (orange) mitotic chromosomes (normalised merged signal for n=4+4). For each locus, gene annotations are shown above; MACS2 consensus ATAC-seq peaks are shown below, with significantly altered peaks in red (DESeq2, padj<0.1). Published CTCF peak locations from asynchronous mouse preB cells^56^ are shown underneath. e. Top ten enriched motifs in peaks which show increased accessibility in *Pbk^−/−^*mitotic chromosomes. C2H2-ZF proteins are indicated in bold. f. Differential footprinting analysis of *Pbk^−/−^* versus *Pbk^+/+^* mitotic chromosome ATAC-seq data in consensus peaks, using TOBIAS with motifs from JASPAR (2024 core vertebrates). Motifs in the top 5% by both p-value and size of change are highlighted in red and associated TFs are annotated (Thap11 label is omited for clarity); bold font indicates C2H2-ZF proteins. g. Nucleosome-free (≤100bp) ATAC-seq signal from *Pbk^+/+^* (middle) and *Pbk^−/−^*(right) mitotic chromosomes at known CTCF binding sites. CTCF peak locations and ChIP-seq coverage (left) are taken from asynchronous primary mouse preB cells^56^. Windows are centred on CTCF motifs, and coverage (10 bp bins) is shown for individual peaks (lower, n=8,532 peaks), ordered by total CTCF signal, with average signal ploted above. h. Average nucleosomal (180-250 bp) ATAC-seq coverage at CTCF peaks, centred on CTCF motifs, for asynchronous WT mouse preB cells (upper^56^) and *Pbk^+/+^* and *Pbk^−/−^* mitotic chromosomes (lower). CTCF ChIP-seq coverage is shown for WT asynchronous cells (upper, black line); doted lines indicate average mid-points of flanking nucleosomes (separation: WT asynchronous=441 bp, *Pbk^+/+^* mitotic chromosomes=399 bp, *Pbk^−/−^* mitotic chromosomes=421 bp). i. Model for how PBK regulates CTCF dissociation, chromatin accessibility and nucleosome repositioning in mitosis. CTCF binds in interphase, maintaining a nucleosome-free accessible region, with well-positioned flanking nucleosomes (upper). In WT mitotic cells (middle, blue), CTCF is evicted by PBK-mediated phosphorylation, leading to an inward shift of nucleosomes and a reduction in accessibility, consistent with previous data and models^13,57^. In cells lacking PBK (lower, orange), CTCF remains unphosphorylated and is at least partially retained on mitotic chromosomes, resulting in greater protection of the nucleosome-free accessible region and a reduced inward shift of flanking nucleosomes.

### CTCF retention on *Pbk^−/−^* mitotic chromosomes leads to increased accessibility and altered nucleosome positioning

In order to probe for evidence of differential DNA binding of individual factors between *Pbk^+/+^* and *Pbk^−/−^* samples, we performed TF footprinting analysis on our ATAC-seq data (Figure 5f). These results revealed increased footprint protection for a wide range of C2H2-ZF proteins, consistent with generally increased retention of these factors on *Pbk^−/−^* mitotic chromosomes. Although significant, most footprint differences were relatively modest; however, in contrast to these subtle differences, CTCF motifs clearly showed enhanced protection in the *Pbk^−/−^* samples (Figure 5f and Supplementary Figure S5f), consistent with this factor remaining chromosome-associated in mitosis (Figure 4d). Focusing on published interphase CTCF-binding sites^56^ (Figure 5g, left), we observed increased chromatin accessibility at these loci in *Pbk^−/−^* compared to *Pbk^+/+^* mitotic chromosomes (Figure 5g), in agreement with the motif enrichment results shown in Figure 5e. Nucleosome positioning around known CTCF-binding sites was also altered on *Pbk^−/−^* mitotic chromosomes (Figure 5h), such that flanking nucleosomes were shifted slightly outwards from CTCF motifs (421 bp spacing) as compared with mitotic *Pbk^+/+^*spacing (399 bp), and more closely resembled the spacing seen in WT asynchronous samples (441 bp). These data are consistent with widespread mitotic dissociation of CTCF in preB cells mediated by PBK phosphorylation, that is effectively hampered in the absence of PBK. Our observations are also in agreement with a previously suggested model whereby mitotic dissociation of CTCF leads to an inward shift of flanking nucleosome positions^57^. Collectively these results suggest that in the absence of PBK, CTCF remains chromosome-associated throughout mitosis and results in a greater preservation of nucleosome-free accessible regions by preventing adjacent nucleosomal array repositioning (Figure 5i).

## Discussion

It has been proposed that transcription factors that remain associated with chromosomes throughout mitosis could ‘bookmark’ the genome for future rapid expression of genes in daughter cells as they enter G1^1,11,58–62^. Mitotic bookmarking factors have been described in many different cell types, including ESRRB in mouse embryonic stem cells (mESCs)^12,63^, GATA1 in erythroblasts^60^, GATA2 in haemopoietic precursors^4^, FOXA1 in hepatocytes^61^ and RUNX2 in osteoblasts^64^. However, we have previously estimated that only approximately ten percent of proteins that were detected in mitotic mESC lysate pellets were enriched on chromosomes at metaphase^8^. Most chromatin-bound factors were presumed to dissociate as a consequence of reduced transcription, altered DNA binding kinetics, physical compaction of chromosomes, or in response to mitotic kinase-mediated phosphorylation and eviction^65^. In addition, alterations to the chromatin environment, for example depletion of H3K9me3, can also alter the retention of specific bookmarking factors on metaphase chromosomes^49^. Here we investigated the mitotic eviction of the lymphoid-specific DNA binding factor Ikaros. We showed that a mitotic kinase PBK (also known as TOPK), that was independently discovered by two groups more than twenty years ago^66,67^ and causes global phosphorylation of C2H2-ZF proteins^45,46^, was both necessary and sufficient to release Ikaros from mitotic chromosomes. Genetic deletion of PBK from mouse preB cells prevented Ikaros displacement during metaphase. Furthermore, in WT cells, Ikaros rapidly re-associated to metaphase chromosomes when treated *in situ* with the PBK inhibitor OTS514, reminiscent of the rapid re-localisation to centromeric regions observed at anaphase/telophase in normal mitosis. Our results therefore show that Ikaros recruitment to chromosomes is regulated by a dynamic interplay between the mitotic-specific activity of PBK and an opposing phosphatase, most likely PP1 which has been shown to regulate Ikaros binding in interphase^51,68,69^. Furthermore, our data from T cells and macrophages suggest that this mechanism is shared between different cell types.

While PBK can mediate the phosphorylation of many C2H2-ZF proteins including SP1 and YY1 (as well as many others exemplified in Figure 4c), it is worth noting that SP1 and YY1 did not show re-localisation to mitotic chromosomes in PBK-null cells. Although ATAC-seq footprinting analysis detected differences at multiple C2H2-ZF motifs, most changes were very subtle, suggesting relatively minor alterations in the occupancy or affinity of these factors in the absence of PBK. This implies that many targets of PBK are also regulated by other kinases which can phosphorylate and displace these factors from mitotic chromosomes, which is consistent with published literature. For example, YY1 is reported to be targeted by Aurora kinases^17^, including inactivation by Aurora kinase A-mediated phosphorylation at zinc finger serine 365^18^. Interestingly, we noted that SP1 shares a highly similar potential Aurora A target motif (Supplementary Figure S6a), whilst DNA-binding of SP1 in mitosis can also be regulated by CDK1^19,20^. In contrast to this, a subset of C2H2-ZF factors, including Ikaros, Aiolos, CTCF, ZNF131 and ZBTB11, which all lack putative Aurora A target motifs (Supplementary Figure S6a), appear to be uniquely dependent on PBK for mitotic displacement. Such distinctions may be important when considering the likely impacts of PBK/TOPK inhibitors in future cancer therapeutics^70^.

Among the C2H2-ZF targets of PBK-mediated phosphorylation, we identified CTCF. In dividing preB cells we show that CTCF is largely displaced from chromosomes by metaphase but remains chromosome-associated in PBK-null cells, demonstrating unequivocally that displacement of CTCF from chromosomes is regulated by PBK. ATAC-seq comparisons made between *Pbk^+/+^* and *Pbk^−/−^*mitotic chromosomes confirm this view, revealing enhanced footprint protection, increased accessibility and altered nucleosome positioning at CTCF binding sites in the absence of PBK. CTCF can also stabilise the binding of cohesin complexes^71,72^, and in this context it is interesting to note that mitotic depletion of cohesin appears less pronounced in *Pbk^−/−^* cells where CTCF is retained. Intriguingly, despite more than a decade of intense investigation, the status of CTCF in interphase^73,74^ and as a mitotic bookmark^10,13,57,62,71,75–78^, remains enigmatic. In interphase, CTCF plays a key role in forming topologically associated domains (TADs) and demarcating chromatin loops^72,74,79–81^, but by prometaphase such structures are no longer detected^57,78,82^ and CTCF is generally thought to largely be dissociated^6,57,83,84^. Several reports have suggested that CTCF can ‘bookmark’ a minority of sites through mitosis, at least in certain cells^13,62,75,76^, and it is proposed that this could render candidate genes more susceptible to rapid reactivation in G1^13,62^. Our data are not necessarily at odds with such claims, but do show that the majority of CTCF is efficiently evicted from preB cell chromosomes by metaphase. Interestingly, prior reports in which a degron was used to remove CTCF during mitosis did not show widespread reduction in the rapid re-expression of proposed CTCF bookmarked genes^62^. It has subsequently been proposed that mitotic bookmarking of the genome could be achieved by conjoint factors that effectively recognise similar or identical binding sites and operate (in tandem) throughout the cell cycle^5^. In this regard, we have previously shown that ADNP and ADNP2 are retained by mitotic mESC chromosomes^8^ (illustrated in Supplementary Figure S6b for reference), and these factors form part of the ChAHP complex that competes with CTCF for a common set of binding motifs^85,86^. Whether there is potential overlap in mitotic binding, and therefore potential redundancy of CTCF and ADNP functions in cell cycle, will be of interest to explore in the future.

Mitotic chromosomes isolated from preB cells that lacked PBK were significantly larger (less compact) than their normal counterparts. This was demonstrated both with purified individual chromosomes 3 and 19 that were stained with Hoechst and Chromomycin dyes and isolated by flow cytometry, or by direct analysis of these chromosomes on metaphase spreads. Although PBK has previously been suggested to contribute to chromosome condensation via H3S10 phosphorylation^55^, we did not observe any reduction in this modification in *Pbk^−/−^* mitotically arrested cells. These observations, together with widespread increases chromatin accessibility, imply that a failure to correctly displace C2H2-ZF binding proteins at prophase/metaphase may contribute to compromised mitotic chromosome compaction and structure, a result which is broadly in line with claims that chromosome condensation in mitosis requires global reductions in transcription, acetylation and transcription factor binding^65,87^. In addition to C2H2-ZF factors, we also observed an apparent increased retention of other chromatin-related proteins, in particular nucleosome remodelling complexes, an observation which could potentially be explained by TF interactions, such as between Ikaros and NuRD^38–41^. The observation that *Pbk^−/−^* chromosomes appear to retain more heterochromatin proteins and constituents of the NuRD deacetylase complex, yet are less compact, may seem counterintuitive. Previously we have shown that several mediators of repressive chromatin, such as DNA methylation and PRC2-mediated histone H3K27me3 remained chromosome-associated throughout mitosis, and their deletion resulted in mitotic chromosomes being less condensed than their normal counterparts^8^. On the other hand, removal of the heterochromatin mark H3K9me3 resulted in mitotic chromosomes which were considerably smaller and more compact than their WT equivalents^49^. In addition, cohesin, a complex which is largely removed from chromosome arms during prophase but is protected at centromeres by shugoshin^54^, has been shown to provoke a widespread de-condensation of metaphase chromosomes when experimentally cleaved *in situ*^8^. Collectively these data caution against overly simplistic predictions of the direct relationship between individual chromatin components and mitotic chromosome condensation and argue that even low levels of chromosome-associated proteins can have important functional and structural consequences for metaphase chromosomes.

Ikaros/Lyf-1 is one of the earliest and best studied regulators of lymphoid identity, that was initially discovered and characterised more than thirty years ago^24–26^. Since then, it has become clear that Ikaros regulates lineage-specific gene activity and silencing across multiple scales and levels of organisation, harnessing partnerships with a plethora of other transcription factors, family members and chromatin-modifier complexes^30–42,88^. Ikaros is essential for the commitment and differentiation of lymphocytes^27,29^ and its de-regulation can drive the onset and progression of B and T cell leukaemias^89–92^. Here we show that the normal displacement of Ikaros (and its homologue Aiolos) from chromosomes entering mitosis is controlled by a single mitotic kinase PBK, and demonstrated that this is both necessary and sufficient for eviction. This mechanism is shared between cell types, and PBK is also implicated in regulating the mitotic behaviour of CTCF and the NuRD complex, as well as ZNF131 and ZBTB11, two factors that have been shown to prevent the aberrant induction of pro-differentiation genes in pluripotent stem cells^93^. While it is perhaps surprising that mitotic displacement of CTCF and lineage specific transcription factors from chromosomes relies solely on a single kinase, this intimate dependency could offer the opportunity for co-ordinate displacement and dynamic rebinding of a vital subset of PBK targets that will ultimately determine the genomic organisation, acetylation, spatial context and expression of lineage-specifying genes early in G1-phase.

## Methods

### Cells and cell culture

Abelson-transformed WT mouse preB cells^8^ and genetically modified derivatives (see below) were cultured in suspension in IMDM medium supplemented with 12% FCS, 2 mM L-glutamine, 2X non-essential amino acids, antibiotics and 50 µM 2-mercaptoethanol. Mouse VL3-3M2 cells were cultured in suspension in IMDM medium supplemented with 10% FCS, 2 mM L-glutamine, 1 mM sodium pyruvate, antibiotics and 50 µM 2-mercaptoethanol. Mouse J774A.1 cells (adherent) were cultured in DMEM medium supplemented with 10% FCS, 1 mM sodium pyruvate and antibiotics, and were detached with trypsin for passaging. All cells were maintained at 37 °C with 5% CO_2_ and split every 2–3 days.

### Metaphase arrest and propidium iodide staining

PreB cells were diluted to approximately 10^6^ cells/ml and arrested by addition of 0.1 μg/ml demecolcine (Sigma-Aldrich, D1925) for 5 h. For propidium iodide (PI) staining, 10^6^ cells were fixed with ice-cold 70% ethanol and stored at −20°C until staining. Fixed cells were washed once with PBS and incubated with PI stain (1X PBS, 0.05 mg/ml PI (Sigma-Aldrich, P4864), 1 mg/ml RNase A, 0.05% IGEPAL CA-630) for 10 min at room temperature (RT) and 20 min on ice. PI signal was acquired in linear mode using a BD FACSymphony A3 flow cytometer and BD FACSDiva Software (v9.1). FlowJo software (v10.8.1) was used to analyse the data and create plots.

### CRISPR/Cas9 generation of knock-in (KI) and knock-out (KO) mouse preB cells

The UCSC genome browser (https://genome.ucsc.edu/) was used to select gRNA sequences (Table 1)^94,95^. Corresponding oligos were annealed, phosphorylated with T4 PNK (NEB) and cloned into the pU6-(BbsI)_CBh-Cas9-T2A-mCherry plasmid^96^ (a gift from Ralf Kuehn, Addgene plasmid #64324) by golden gate assembly using BbsI-HF and T4 ligase (both NEB). A donor plasmid for *Ikzf1-mNeonGreen* KI was generated by incorporating the *mNeonGreen* sequence, flanked by left and right *Ikzf1* homology arms, into a backbone containing EBFP2 (details in Table 2). The NEBuilder Assembly Tool was used to design primers for amplification of each component of the donor plasmid (Table 2) using Phusion (NEB); purified PCR products were assembled using NEBuilder HiFi DNA Assembly Master Mix (NEB). Assembled gRNA/Cas9 and donor plasmids were transformed into DH5α bacteria and final purified plasmids were checked by sanger sequencing (Genewiz).

**Table 1.**
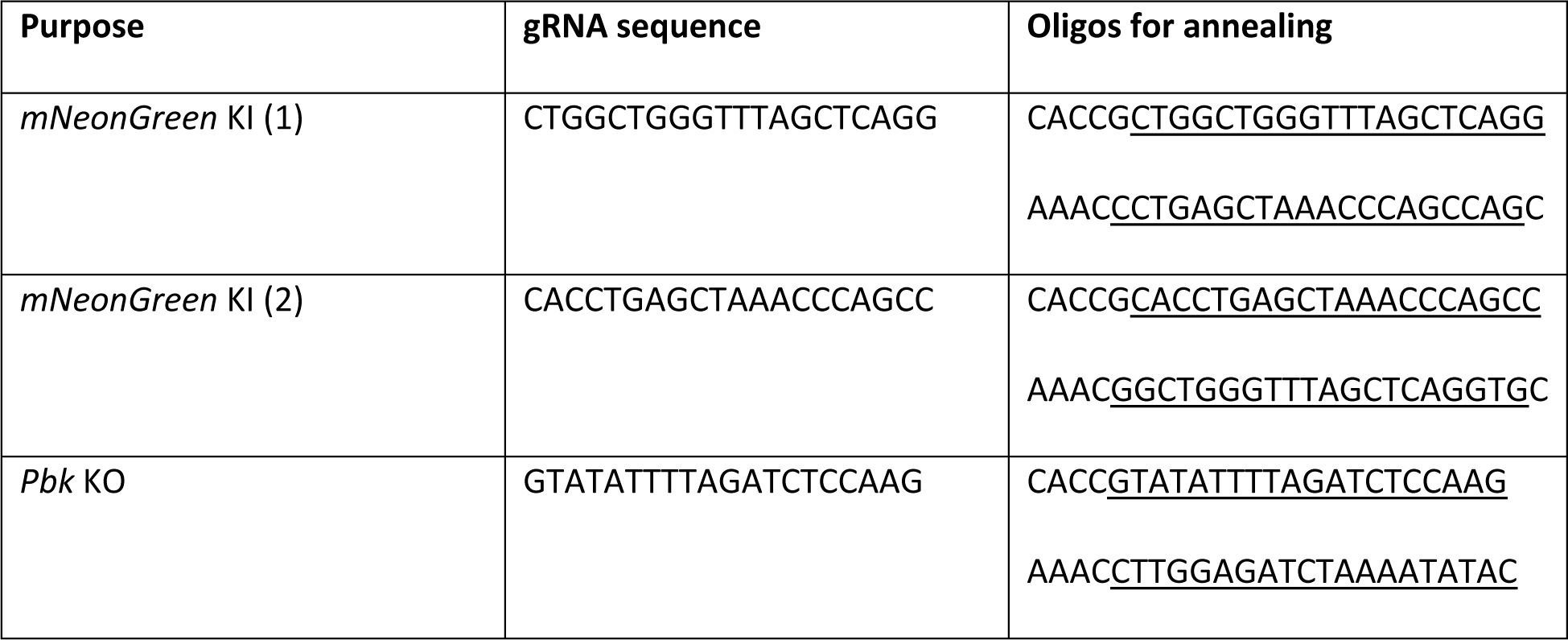
| Guide RNA sequences and corresponding oligos. The UCSC genome browser (https://genome.ucsc.edu/) was used to select gRNA sequences^94,95^ at the 3’ end of *Ikzf1* (two gRNAs selected) or the third coding exon of *Pbk*. Corresponding oligos contain the gRNA sequence (underlined), an upstream “G” if not present in the gRNA sequence (to increase efficiency of U6-driven transcription), and overhangs for golden gate assembly.

**Table 2.**
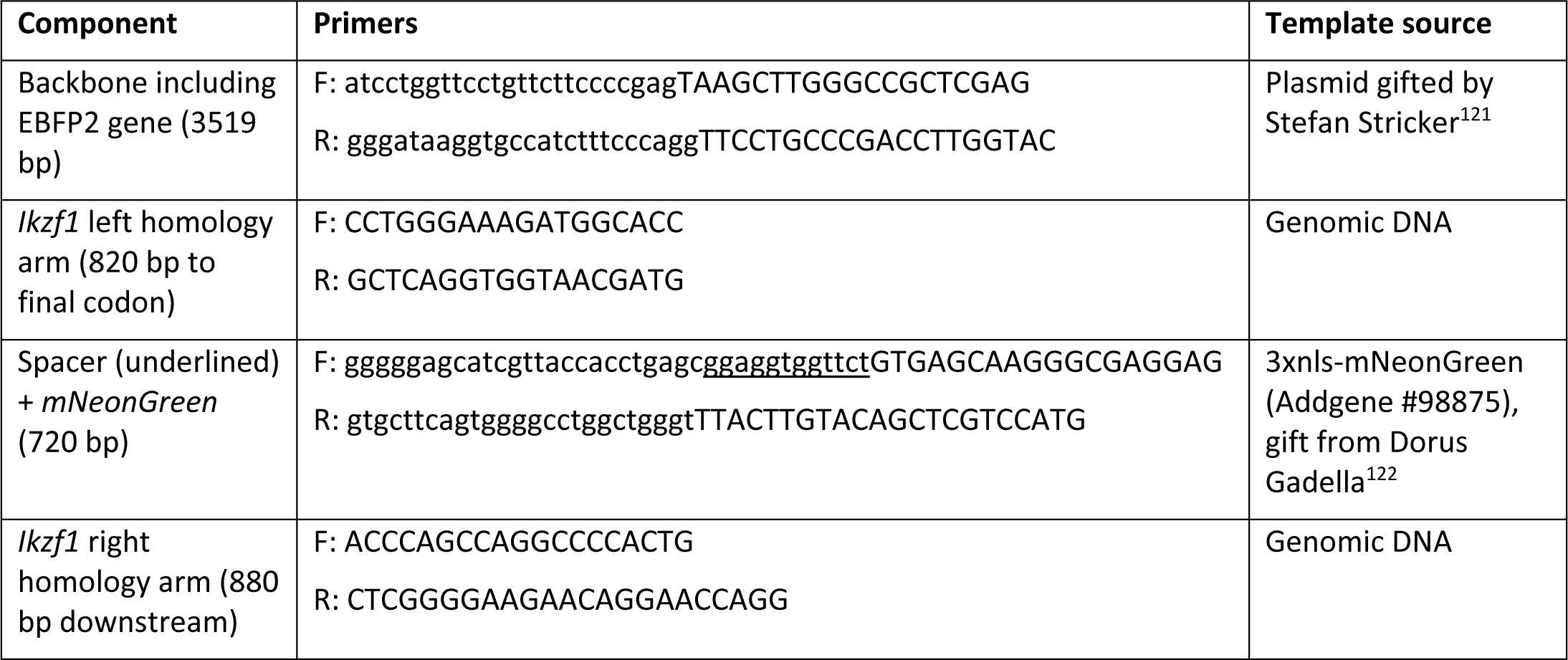
| Components and primers for *Ikzf1-mNeonGreen* donor plasmid construction. Primers were designed using Primer3Plus (https://www.primer3plus.com/, v3.1.0)^120^ and the NEBuilder Assembly Tool (https://nebuilder.neb.com, v2.3.0). Upper case leters anneal to the template, lower case leters provide overlaps for assembly.

PreB cells were transfected with the appropriate gRNA/Cas9 plasmid alone (KO), or in combination with the *mNeonGreen* donor plasmid (KI), using Lipofectamine 3000 (Invitrogen). After 24 h mCherry positive or mCherry/EBFP double positive transfected cells were isolated by FACs (note that efficiency was very low) and cultured as a pool for 1-2 weeks before sorting single cells into 96 well plates. For *mNeonGreen* KI, cells which were positive for mNeonGreen and negative for both mCherry and EBFP were selected. For *Pbk* KO, mCherry negative cells were selected. Clones were grown and screened by PCR (primers in Table 3) and/or western blotting. Genetic modifications were verified by sanger sequencing of PCR products (Genewiz), and clones selected for downstream experiments were confirmed to be karyotypically normal.

**Table 3.**
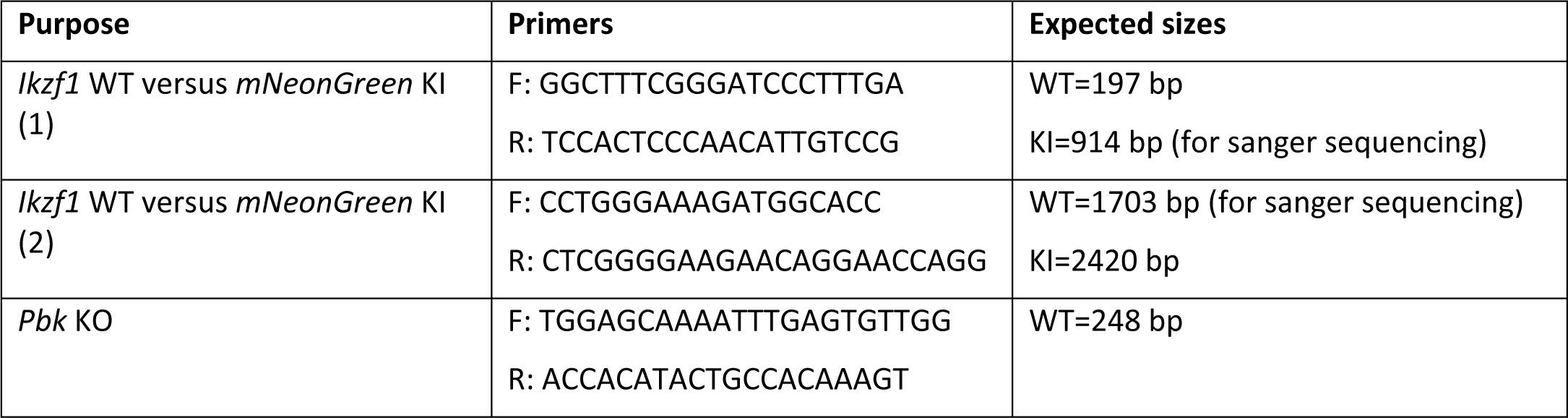
| Primers used for verification of KI or KO cells. Primers were designed using Primer3Plus (https://www.primer3plus.com/, v3.1.0 and v3.2.0)^120^.

### Immunofluorescence (IF)

PreB and VL3-3M2 suspension cells were spun onto poly-D-lysine coated coverslips or µ-Slide 18-well chamber slides (ibidi) whilst J774A.1 adherent cells were grown directly on µ-Slide 18-well chamber slides. For inhibitor treatments, inhibitor-containing medium was added to the cells for 10 min at 37 °C immediately prior to fixation. Cells were fixed with 2% formaldehyde for 15 min at RT, permeabilised with 0.1-0.5% Triton X-100 for 15 min at RT and blocked with 2% bovine serum albumin and 5% goat serum. Primary antibodies (Table 4) were diluted in blocking buffer and incubated with cells overnight at 4 °C. Cells were washed three times with PBS and incubated with goat anti-rabbit secondary antibody (Alexa Fluor 633, Invitrogen, A-21070) diluted 1:500 in blocking solution for 1 h at RT. Cells were washed with PBS and either stained for 5 min with 1 µg/ml DAPI (for chamber slides) or mounted using DAPI-containing Vectashield (Vector Laboratories; for coverslips). An Olympus IX70 inverted microscope with a UPlanApo 100X/1.35 oil iris objective was used for image acquisition with Micro-Manager software (v2.0). Z-stacks were collected with 0.35 µm intervals and images were deconvolved using Huygens Professional software (Scientific Volume Imaging, v19.10), using the CMLE algorithm and default parameters for widefield deconvolution.

**Table 4.**
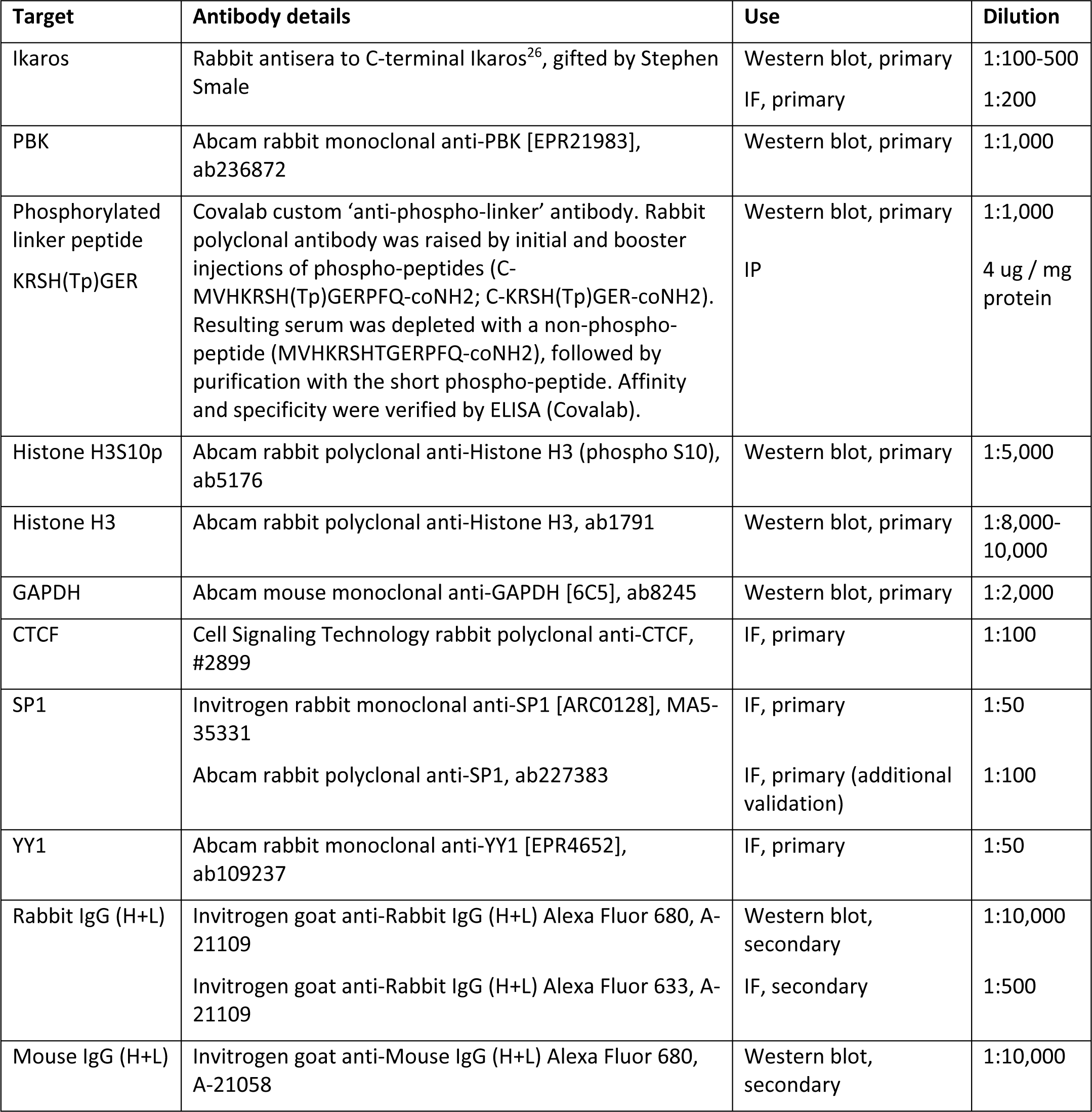
| Antibodies used in this study.

### Live-cell imaging

Ikaros-mNeonGreen KI mouse preB cells were switched to fully supplemented phenol-free medium containing 20 mM HEPES and 0.5-1 µM SiR-DNA (Spirochrome) at least 30 min prior to imaging. Cells were transferred to µ-Slide 8- or 18-well chamber slides (ibidi) and allowed to settle or were briefly centrifuged at 300 *g* before imaging. For testing different inhibitor treatments (Table 5), inhibitors were added immediately prior to transferring cells to chamber slides, and snapshots were collected in a window between 5 min and 15 min after inhibitor addition (average 10 min). For timelapse imaging of individual cells before and after OTS514 addition, a cell in metaphase was identified and imaged. The medium was then carefully removed and replaced with OTS514-containing medium and, after checking the focus, another snapshot was acquired immediately, and at 1 min intervals thereafter. Live-cell snapshots were collected on an Olympus IX70 inverted microscope with a 37 °C environmental chamber and 5% CO_2_ supply using a UPlanApo 100X/1.35 oil iris objective and Micro-Manager software (v2.0). Z-stacks were collected at 0.35 µm intervals and images were deconvolved as for IF. Following deconvolution, volume analysis was performed using the surfaces and cells packages in Imaris (Bitplane, v10.0.0), and mean intensity measurements were performed in both chromosomal and cytoplasmic compartments. Statistics were exported from Imaris, collated using a python script, and the chromosomal to cytoplasmic intensity ratio was calculated.

**Table 5.** | Kinase and phosphatase inhibitors used in this study.

| Inhibitor | Source(s) | Main target(s) | Concentration |
| --- | --- | --- | --- |
| K252a | Abcam, ab120419<br>Enzo Life Sciences (BioVision), 2013-1000 | Broad, non-selective protein kinase inhibitor | 1 μM |
| OTS514 hydrochloride | Stratech (Selleck Chemicals), S7652-SEL | PBK (also known as TOPK) | 10 μM |
| VX-680 (also known as MK-0457 or Tozasertib) | Stratech (ApexBio), A4111-APE | Aurora kinases (and wide array of other kinases) | 10 μM |
| Alsterpaullone (Alp) | Stratech (ApexBio), B7855-APE | Cyclin dependent kinases (CDKs), GSK3β | 10 μM |
| CX-4945 (also known as Silmitasertib) | Stratech (ApexBio), A8330-APE | Casein kinase 2 (CK2) | 10 μM |
| KN-62 | Stratech (Selleck Chemicals), S7422-SEL | CaMKII (also CaMKI, P2RX7, CaMKIV) | 10 μM |
| Chelerythrine Chloride | Stratech (ApexBio), A3306-APE | Protein kinase C (PKC) | 10 μM |
| Akt Inhibitor VIII | Merck (Calbiochem), 124017 | Akt1/2 | 10 μM |
| Saracatinib (also known as AZD0530) | Stratech (ApexBio), A2133-APE | Src family kinases (also Abl and EGFR) | 10 μM |
| Hesperadin | Merck (Calbiochem), 375680 | Aurora kinase B | 10 μM |
| OTS964 | Stratech (Selleck Chemicals), S7648-SEL | PBK (also known as TOPK), CDK11B | 10 μM |
| Okadaic acid (OA) | Merck, O9381<br>Cell Signaling, 5934S | Protein phosphatase PP2A (also PP1 at lower potency) | 1 μM |
| Calyculin A | Merck (Calbiochem), 5082260001<br>Abcam, ab141784 | Protein phosphatases PP1 and PP2A | 0.1 μM |

Time-lapse images for Supplementary Videos S1-4 were collected using an Olympus IX83 microscope equipped with a Yokogawa CSU-W1 spinning disk and Hamamatsu ORCA-Flash 4.0 camera, with a 37°C environmental chamber and 5% CO_2_ supply, using a UPlanSApo 60x/1.35 oil objective. Z-stacks (3 µm step size) were acquired with cellSens Dimension software (v2.3) every 3 min, with autofocus enabled and laser power set at 2%.

### Western blotting

Asynchronous or arrested preB cells were pelleted at 300 *g* and either processed immediately, or snap-frozen and stored at -80 °C. For OTS514 treatment, 10 µM of inhibitor was added to arrested cells for 10 min at 37 °C, before pelleting cells and processing immediately. For processing, cells were resuspended in cold RIPA buffer (50 mM Tris-HCl pH 8.8, 150 mM NaCl, 1% Triton X-100, 0.5% sodium deoxycholate, 0.1% SDS, 1 mM EDTA, 3 mM MgCl_2_) containing 250 U/ml Benzonase (Sigma, E1014), 1X cOmplete EDTA-free protease inhibitor cocktail (Roche, 11873580001), and, for detection of phosphorylated proteins, 1X PhosSTOP phosphatase inhibitor (Roche, 4906845001). Samples were incubated at RT for 20 min before quantification with the Bio-Rad DC protein assay kit. Samples were diluted with RIPA buffer, if required, and mixed 3:1 with 4X Laemmli buffer (250 mM Tris-HCl pH 6.8, 8% SDS, 40% glycerol, 20% β-mercaptoethanol, bromophenol blue). Proteins (15-30 µg) were resolved on a 4-15% or 10% acrylamide gel and transferred onto a PVDF membrane (Invitrogen iBlot 2 or Bio-Rad Trans-Blot Turbo system). Membranes were blocked with 5% milk (standard protein detection) or 5% BSA (phosphorylated protein detection) in TBS-T (TBS with 0.1% Tween 20) for 1 h at RT before incubating with primary antibodies (Table 4) overnight in blocking solution. After three washes in TBS-T, membranes were incubated with appropriate secondary antibody (Table 4) for 1 h at RT in blocking solution. Following three washes in TBS-T, fluorescence imaging of the membrane was performed with the LI-COR Odyssey CLx system using Image Studio software (v5.2.5).

### Chromosome size measurements from metaphase spreads

Arrested preB cells were lysed in hypotonic solution (75 mM KCl, 10 mM MgSO_4_, pH 8) for 25 min at 37 °C, before centrifuging at 500 *g* for 8 min to pellet nuclei. Nuclei were resuspended in residual supernatant and stored in fixative (75% methanol, 25% acetic acid) at -20 °C. To prepare metaphase spreads, nuclei were pelleted (500 *g*, 8 min), washed three times in fixative, and resuspended in a small volume of fixative (to a pale grey appearance). Next, 23 μl of sample was dropped directly onto a 20 μl drop of 45% acetic acid on a glass Twinfrost slide, tilting to spread nuclei, before air-drying slides at RT.

Chromosomes 3 and 19 were detected with mouse chromosome painting probes (Metasystems Probes) according to the manufacturer’s protocol before samples were stained with DAPI (1 µg/ml, 5 min) and mounted in Vectashield (Vector Laboratories). Images were collected with a Leica SP5 II confocal microscope using LAS-AF software (v2.7.3.9723). Chromosome spread images were first segmented using Cellpose (v2.0)^97^ with the ’cyto’ model, with additional training performed to improve the segmentation precision of individual chromosomes in clumped spreads. Labelled mask images were subsequently imported into QuPath (v0.4.3)^98^ for chromosome paint classification and area measurements.

### Isolation of mitotic chromosomes by flow cytometry

Mitotic chromosomes were purified by flow cytometry as previously described^8,49^, with minor modifications. Briefly, arrested preB cells (∼10^8^) were incubated for 20 min at RT in 10 ml of hypotonic solution (75 mM KCl, 10 mM MgSO4, 0.5 mM spermidine trihydrochloride (Sigma-Aldrich, S2501), 0.2 mM spermine tetrahydrochloride (Sigma-Aldrich, S2876), pH 8.0), followed by 15 min on ice in 3 ml of polyamine buffer (80 mM KCl, 15 mM Tris-HCl, 2 mM EDTA, 0.5 mM EGTA, 3 mM DTT, 0.25% Triton X-100, 0.5 mM spermidine trihydrochloride, 0.2 mM spermine tetrahydrochloride, pH 7.7). Samples were vortexed for 30 s at maximum speed, passed several times through a 21-gauge needle, centrifuged at 200 *g* for 2 min, filtered through a 20 μm CellTrics filter (Sysmex) and stored at 4 °C overnight. Next day, chromosomes were stained on ice by adding 5 μg/ml Hoechst 33258 (Sigma-Aldrich, 94403), 25 μg/ml Chromomycin A3 (Sigma-Aldrich, C2659) and MgSO_4_ (10 mM final) for 45 min, followed by addition of sodium citrate (10 mM final) and sodium sulfite (25 mM final) for 1 h.

Stained chromosomes were purified as reported^8,49^, using a Becton Dickinson Influx with BD FACS Sortware (v1.2.0.142), using a 70 μm nozzle tip, a drop-drive frequency of 96 kHz and a sheath pressure of 448 kPa. Forward scatter was measured with a 488 nm laser (Coherent Sapphire, 200 mW); Hoechst 33258 was analysed with a 355 nm air-cooled laser (Spectra Physics Vanguard, 350 mW) and 400 nm (long-pass)/500 nm (short-pass) filters; Chromomycin A3 was analysed with a water-cooled 460 nm laser (Coherent Genesis, 500 mW) and 500 nm (long-pass)/600 nm (short-pass) filters.

### Size measurements of flow-sorted chromosomes

Purified chromosomes 3 or 19 were cytocentrifuged (Cytospin3, Shandon) at 163 *g* for 10 min onto Polysine slides (VWR). After mounting with Vectashield containing DAPI (Vector Laboratories), chromosomes in the centre of the slide were imaged on an Olympus IX70 inverted microscope using a UPlanApo 100X/1.35 oil iris objective and Micro-Manager software (v2.0). Chromosome size analysis was performed in Fiji (1.54e)^99^: following background subtraction and median filtering, images were automatically thresholded and binarised for individual chromosome segmentation and subsequent area measurements. Images were manually inspected to exclude mis-identified chromosomes or centromeres.

### LC-MS/MS analysis of mitotic chromosome samples

Flow-purified chromosomes (10^7^) or unpurified chromosomes from the pre-sorted lysate (85-100 μl) were pelleted at 10,000 *g* for 10 min at 4 °C; pellets were snap-frozen and stored at -80 °C. Pellets were processed by trypsin digest using the iST Sample Preparation Kit (PreOmics, P.O.00001), with minor modifications to the manufacturer’s protocol as follows: in step 1.1, 30 μl lysis buffer was used and samples were sonicated in an ultrasonic bath (1 min) both before and after heating; in steps 3.8-9, samples were resuspended in 20 μl and subjected to both sonication and shaking.

LC-MS/MS analysis was performed using an UltiMate 3000 RSLC nano-flow liquid chromatography system (Thermo Scientific) coupled to a Q-Exactive HF-X mass spectrometer (Thermo Scientific) via an EASY-Spray source (Thermo Scientific) as previously reported^8^, with minor variations as follows. Samples were loaded onto a trap column (Acclaim PepMap 100 C18, 100 μm x 150 mm) at 10 μl/min in 2% acetonitrile and 0.1% trifluoroacetic acid, eluted to an analytical column (Acclaim Pepmap 100 C18, 75 μm x 50 cm) at 250 nl/min, and separated using an increasing gradient of buffer B (75% acetonitrile, 5% DMSO, 0.1% formic acid) in buffer A (5% DMSO, 0.1% formic acid): 1-5% for 5 min, 5-22% for 70 min, 22-42% for 20 min. Eluted peptides were analysed by data-dependent acquisition as described^8^, except with maximum injection time for MS2 of 110 ms. For quality control purposes, pooled replicate samples (pre-sorted or purified chromosome replicates were pooled separately) were run at the start, middle and end to verify consistency throughout the run.

Raw data were processed with MaxQuant (v1.6.10.43)^100^, using the in-built Andromeda search engine against the UniProt *Mus musculus* one-gene-per-protein database (v20230131; 21,976 entries) and against a universal protein contaminants database^101^ (downloaded 20220604; 381 entries). A reverse decoy search approach was used at a 1% false discovery rate (FDR) for both peptide spectrum matches and protein groups (parameters included: max missed cleavages=3; fixed modifications=cysteine carbamidomethylation; variable modifications=methionine oxidation, protein N-terminal acetylation, asparagine deamidation, cyclisation of glutamine to pyro-glutamate). Label-free quantification (LFQ) was performed (LFQ min ratio count=2) and ‘match between runs’ was enabled (match time limit=0.7 min, alignment time limit=20 min). Two pre-sorted lysate pellet samples (*Pbk^+/+^* replicate 1 and *Pbk^−/−^* replicate 4) exhibited outlier retention time distributions and were excluded from the analysis.

Subsequent data analyses were carried out in Perseus (v1.6.15.0)^102^, comparing purified chromosomes (n=4) to pre-sorted lysate pellet (n=3, due to excluded samples) for each of *Pbk^+/+^* and *Pbk^−/−^*conditions to assess chromosomal enrichment or depletion of factors. Proteins with ‘only identified by site’ or ‘reverse’ annotations were removed, and data were log2 transformed. Pearson correlation values (LFQ intensities) were extracted using the multi scatter plot function in Perseus and visualised as a heatmap using the R package corrplot (v0.92). Data were filtered in Perseus to keep proteins with ≥3 valid replicate LFQ intensities per experimental group (purified chromosomes or pre-sorted lysate pellet); the volcano plot function was used to perform a t-test (modified two-tailed t-test: permutation-based FDR<0.05, randomisations=250, S0=0.1) and for subsequent visualisations. Subsets of proteins were highlighted on volcano plots according to GO term and PROSITE annotations assigned in Perseus (mainAnnot.mus_musculus.txt), with additional manual curation using the MGI Gene Ontology Browser (https://www.informatics.jax.org/function.shtml) where required. The following definitions were used: Histones=gene name contains “Hist” or “H2a” (15 in *Pbk^+/+^*; 14 in *Pbk^−/−^*); Proteasome=GOCC name contains “proteasome” (32 in *Pbk^+/+^*; 36 in *Pbk^−/−^*); Cohesin=GOCC name “Cohesin complex” plus Stag1/2 (8 in *Pbk^+/+^*; 7 in *Pbk^−/−^*); Condensin=GOCC name “Condensin complex” or protein name contains “condensin” (8 in *Pbk^+/+^*; 8 in *Pbk^−/−^*); C2H2-Zinc Finger=PROSITE PS00028 or PS50157 (43 in *Pbk^+/+^*; 52 in *Pbk^−/−^*); NuRD Complex=GOCC name “NuRD complex” plus Mbd2 and GATAD2b (11 in *Pbk^+/+^*; 12 in *Pbk^−/−^*).

GO term overrepresentation analysis was performed using PANTHER^103^ (https://www.pantherdb.org/; v18.0; Fisher’s exact test with FDR correction; GO Ontology database DOI: 10.5281/zenodo.8436609; GO cellular component complete). Analysis was performed for factors enriched or depleted on mitotic chromosomes (run together), for each of *Pbk^+/+^* and *Pbk^−/−^*conditions (run separately), using the total proteins detected in each condition as background (all proteins displayed on respective volcano plots). The first majority protein ID per hit was used as input to PANTHER, with unassigned IDs manually updated using the second ID or gene name where possible (*Pbk^+/+^* mapped IDs: background=2,007, enriched=341, depleted=503; *Pbk^−/−^* mapped IDs: background=2,281, enriched=400, depleted=679). GO terms were filtered for significance (FDR<0.05) in at least one condition, and the top most overrepresented terms in either condition were selected for visualisation (14 terms each for mitotically enriched or depleted factors), ordered by the difference in fold overrepresentation between *Pbk^+/+^* and *Pbk^−/−^* conditions.

### Immunoprecipitation (IP)

Arrested preB cells (∼3 x 10^7^) were washed twice in ice-cold PBS, snap-frozen and stored at -80 °C until processing. Cells were lysed at RT for 10 min in 1 ml RIPA buffer containing 250 U/ml Benzonase (Sigma), 1X cOmplete EDTA-free protease inhibitor cocktail (Roche), and 1X PhosSTOP phosphatase inhibitor (Roche) before centrifuging at 16,000 *g* for 10 min at 4 °C. Mitotic lysates were quantified with the Bio-Rad DC protein assay kit and diluted to 0.8 mg/ml with RIPA buffer containing protease and phosphatase inhibitors.

For IP, 1 ml (0.8 mg) of diluted mitotic lysate (input) was incubated overnight with 3.2 µg anti-phospho-linker antibody (Table 4) at 4 °C with end-to-end rotation. MagReSyn Protein G beads (ReSyn Biosciences, MR-PRG002) were washed three times in RIPA buffer, and 10 µl of resuspended beads were added to each IP sample and incubated for 10 min at RT, followed by 1 h at 4 °C, with rotation, to capture antibody-antigen complexes. Using a magnetic separator, the supernatant was removed, and the beads were washed five times with RIPA buffer (first wash with protease/phosphatase inhibitors) and three times with 20 mM EPPS, each wash for 5 min at 4 °C, with rotation. In the final wash, samples were transferred to 0.2 ml PCR tubes, all supernatant was removed, and beads were snap-frozen and stored at -80 °C until processing for mass spectrometry analysis.

### LC-MS/MS analysis of IP and input samples

#### IP sample preparation

Dry beads underwent on-bead digestion in 50 µl digestion solution (20 ng/μl trypsin (Pierce™, 90059), 10 ng/μl lysyl (WAKO, 129-02541), 50 mM EPPS) for 2 h at 37°C, followed by removal of solution from the beads and overnight digestion at RT. The next day, desalting was performed using 100 g/l Oasis HLB beads (Waters, 186007549; 100 μg beads to 1 μg peptides) distributed in an OF1100 orochem filter plate, and a vacuum manifold. Beads were washed twice with 100 μl acetonitrile and three times with 100 μl 0.1% trifluoroacetic acid (TFA). Samples were acidified to 0.5% TFA, loaded onto a plate, washed twice with 400 μl 0.1% TFA, once with 200 μl H_2_O, centrifuged at 1,000 *g* and eluted in 20 μl 70% acetonitrile. Samples were dried by vacuum centrifugation and resuspended in 30 μl 0.1% TFA for mass spectrometry analysis.

#### Input (mitotic lysate) sample preparation

Samples were prepared for Protein Aggregation Capture^104,105^ by mixing samples with SDS (1% final concentration), EPPS pH 8.5 (50 mM final concentration), chloroacetamide (20 mM final concentration) and TCEP (tris(2-carboxyethyl)phosphine, 10 mM final concentration). After heating at 60 °C for 5 min and leaving on the bench for 30 min, prewashed hydroxyl beads (ReSyn Biosciences, MR-HYX010) were added at a protein-to-bead ratio of 1:5. Ethanol was added to a final concentration of 70%, pipetted up and down, left for 5 min to observe bead aggregation, and washed three times with 100 μl 80% ethanol. For digestion, 10 ng/μl trypsin and 5 ng/μl LysC in 100 mM ammonium bicarbonate pH 8 were added and the mixture was incubated for 18 h at 37°C with shaking at 1,600 rpm.

#### LC-MS/MS analysis

Chromatographic separation was performed using an UltiMate 3000 RSLC nano-flow liquid chromatography system (Thermo Scientific) coupled to a Q-Exactive mass spectrometer (Thermo Scientific) via an EASY-Spray source (Thermo Scientific). Electro-spray nebulisation was achieved by interfacing to Bruker PepSep emitters (10 µm, PSFSELJ10). Peptide solutions were injected directly onto the analytical column (self-packed column, CSH C18 1.7 µm beads, 300 μm x 30 cm) at a working flow rate of 5 μl/min for 4 min. Peptides were then separated using a 120 min stepped gradient: 0-45% of buffer B (75% MeCN, 5% DMSO, 0.1% formic acid) in buffer A (5% DMSO, 0.1% formic acid), followed by column conditioning and equilibration. Eluted peptides were analysed by the mass spectrometer in positive polarity, using a data-dependent acquisition mode. An initial MS1 scan (resolution=140,000; AGC target=3e6; maximum injection time=50 ms; range=400-1800 m/z) was followed by eight MS2 scans (analytes with +1 and unassigned charge state excluded; resolution=35,000; AGC target=1e5, maximum injection time=128 ms; intensity threshold=2e3). Normalised collision energy was set to 27%, dynamic exclusion was set to 45 s, total acquisition time was 150 min.

#### Data processing

Raw data were processed with MaxQuant (v1.6.10.43)^100^, searching against the UniProt *Mus musculus* one-gene-per-protein database (v20220720; 21,992 entries), using the same parameters as for chromosome LC-MS/MS samples (see above). Label-free quantification was performed (LFQ min ratio count=1) and ‘match between runs’ was enabled (match time limit=0.7 min, alignment time limit=20 min).

Subsequent data analyses were carried out in Perseus (v1.6.15.0)^102^, comparing *Pbk^+/+^* versus *Pbk^−/−^*input or *Pbk^+/+^* versus *Pbk^−/−^* IP. Proteins with ‘only identified by site’, ‘reverse’ or ‘potential contaminants’ annotations were removed. Proteins were considered robustly detected in only a single condition (*Pbk^+/+^* or *Pbk^−/−^*) if LFQ>0 for all three replicates of one condition and LFQ>0 for maximum one replicate of the other condition with a value ≥2-fold below the mean of the first condition. Proteins detected in at least five samples (across both conditions) were considered significantly different on the basis of a t-test (modified two-tailed t-test: permutation-based FDR<0.05, randomisations=250, S0=0.1). Candidate PBK targets were defined as those hits which were robustly detected after IP from *Pbk^+/+^*but not *Pbk^−/−^* lysates, or which were significantly lower after IP from *Pbk^−/−^* lysates. There was no overlap between these candidates and those proteins which were significantly higher or only detected in *Pbk^+/+^*compared to *Pbk^−/−^* input samples. Normalised LFQ values (fold over max) for candidate PBK targets were displayed as a heatmap using conditional formatting in Microsoft Excel. PROSITE annotations were assigned in Perseus (mainAnnot.mus_musculus.txt) by UniProt majority protein IDs, with missing annotations supplemented manually from UniProt (https://www.uniprot.org/). C2H2-ZF proteins were defined by matches to PROSITE accession numbers PS00028 and/or PS50157.

### ATAC-seq libraries

ATAC-seq was performed on mitotic chromosomes^8,49^, using chromosomes purified from the same preparations as the proteomics analysis (n=4). Briefly, 2 x 10^6^ flow-sorted chromosomes were pelleted at 10,000 *g* for 10 min at 4 °C before tagmentation in a 50 μl reaction (2.5 μl Tagmentase (loaded Tn5 transposase, Diagenode, C01070012), 25 μl 2X Tagmentation buffer (Diagenode, C01019043), 22.5 μl H_2_O) at 37 °C for 30 min with 1,000 rpm shaking. DNA was purified (Qiagen MinElute PCR purification kit) and PCR amplified (7 cycles; NEBNext High Fidelity master mix; primer sequences in Table 6). Amplified libraries were purified using 1.8X volume Ampure XP beads (Beckman Coulter), firstly using 0.5X beads to remove large fragments, before using the remaining volume of beads to capture and purify libraries. Purified ATAC-seq libraries were assessed by Bioanalyzer and quantified by Qubit and KAPA Library Quantification (Roche). Paired-end sequencing (60 bp) was carried out on an Illumina NextSeq 2000 (NextSeq 1000/2000 Control Software v1.4.1.39716; primary analysis RTA 3.9.25; secondary analysis DRAGEN Generate FastQ v3.7.4; reads demultiplexed with bcl2fastq2 v2.20 (allowing 0 mismatches)).

**Table 6.**
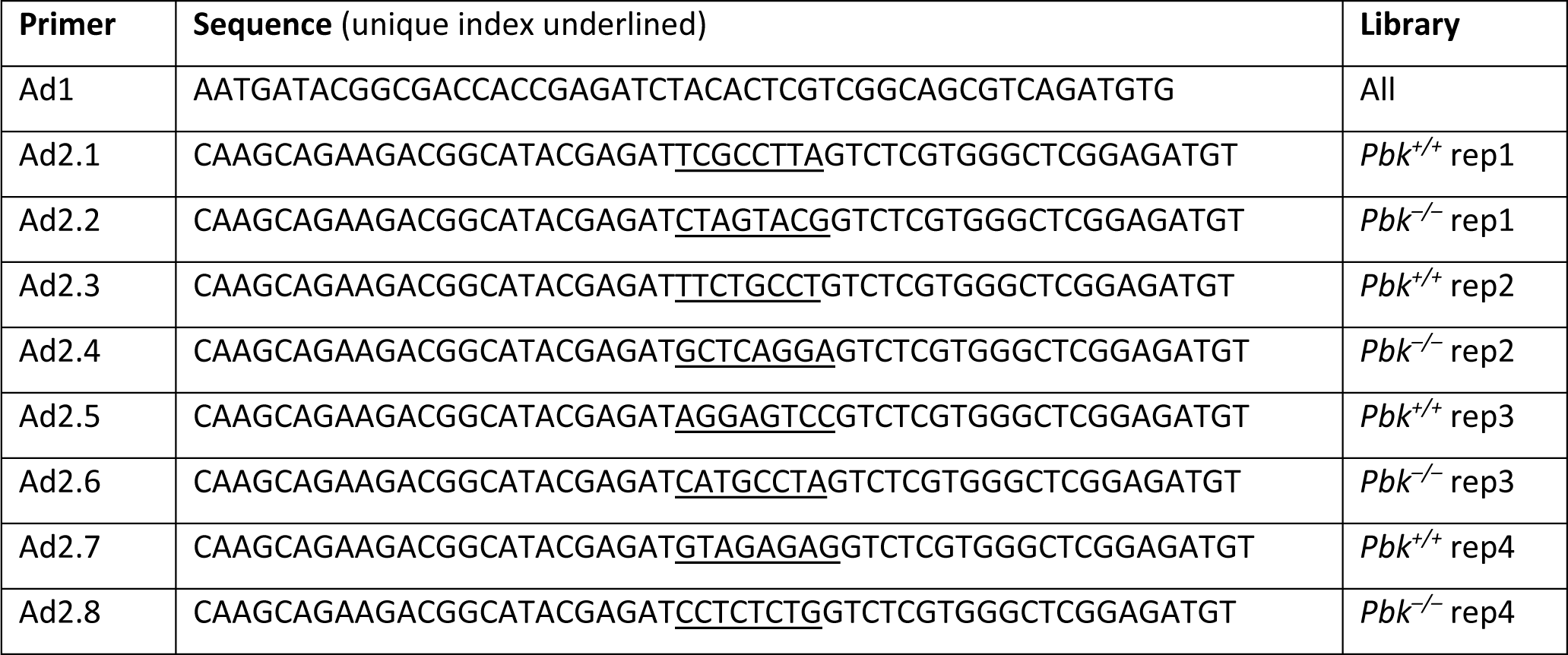
| Primers for ATAC-seq library amplification. Primer sequences were taken from^123^ and ordered with HPLC purification (Sigma-Aldrich).

### Sequencing data analysis

#### ATAC-seq data processing

ATAC-seq reads from sorted chromosomes were trimmed with fastp (v0.23.3)^106^ to remove adapters and nucleotides with quality <20. Reads were aligned to GRCm39 with bwa-mem (v0.7.17)^107^ and filtered using samtools (v1.17)^108^ to retain only properly paired alignments with a quality score of ≥3. PCR duplicates were marked using sambamba (v1.0.1)^109^ and replicates were merged using samtools (v1.17). Fragment size distributions were calculated for merged replicates using deeptools bamPEFragmentSize (v3.5.1)^110^. The fragment size distributions shown in Supplementary Figure S5a are smoothed for better visualisation by taking a running mean in 5 bp windows.

Peaks were called using MACS2 callpeak (v2.2.8)^111^ with the parameters “--nomodel --shift -100 --extsize 200 -g mm” to call accessible regions for each replicate individually and for merged replicates. “Consensus” peak sets for *Pbk^+/+^* and *Pbk^−/−^* were created by taking peaks called in the merged replicates that were also called in at least 3 out of 4 individual replicates. Peaks and data on chromosomes 6 and 14 were excluded from downstream analysis because by visual inspection these chromosomes have globally reduced ATAC-seq coverage in *Pbk^−/−^*.

#### Differential accessibility

For differential accessibility analysis, ATAC-seq read alignments were shifted to reflect the Tn5 cut site positions (+4 bp for plus strand, -5 bp for minus strand) and resized to 1 bp, using functions from the GenomicRanges R package (v1.46.1)^112^. Consensus accessible peaks from *Pbk^+/+^* and *Pbk^−/−^* were merged and cut sites overlapping each merged peak in each replicate were counted. The counts were used as input for differential accessibility analysis with DESeq2 (v1.34.0)^113^. Shrunken log2 fold changes (using type=“apeglm”) were visualised uisng an MA plot (Figure 5b) and the variance stabilizing transformation was applied before performing principal component analysis (Supplementary Figure S5b). Peaks with an adjusted p value <0.1 were defined as significantly different between *Pbk^+/+^* and *Pbk^−/−^*.

Motif enrichment analysis in differentially accessible peaks was performed with AME (MEME suite v5.5.4)^114^ using JASPAR 2024 CORE vertebrates non-redundant motifs^115^ and merged consensus peaks from *Pbk^+/+^* and *Pbk^−/−^* as background. GO term enrichment analysis (“Biological Process” terms only) was carried out using clusterProfiler (v4.2.2)^116^. Genes with an annotated TSS overlapping a significantly differential peak were compared to genes with an annotated TSS overlapping any of the peaks used as input to the DESeq2 analysis.

#### Footprinting analysis

Transcription factor footprinting analysis was carried out using TOBIAS (v0.16.0)^117^ with JASPAR 2024 CORE vertebrates non-redundant motifs^115^. Motifs in the top 5% by both p-value and absolute log2 fold change were selected as significant. Footprint plots in Supplementary Figure S5f show adjusted cut site density from TOBIAS in a ±60 bp window around motif centres, with signal smoothed by taking the running mean in 5 bp windows to improve visualisation.

#### Processing of published CTCF ChIP-seq and ATAC-seq data

CTCF ChIP-seq data from preB cells from young mice^56^ were download from SRA (fasterq-dump, sratools v3.0.3) and aligned to GRCm39 using bowtie2 (v2.5.1)^118^ in local mode and filtered using samtools (v1.16.1)^108^ to retain only properly paired alignments with a quality score of ≥30. PCR duplicates were marked using sambamba (v1.0.1)^109^. Peaks were called using MACS2 (v2.2.8)^111^. Very few peaks were called for Rep2, therefore only peaks from Rep1 were used for downstream analysis. CTCF motif locations were identified using the motifmatchr R package (v1.16.0, https://bioconductor.org/packages/motifmatchr) and the MA0139.2 motif from JASPAR2024^115^. Out of 10,424 CTCF peaks, 8,532 had a CTCF motif match.

ATAC-seq data from preB cells from young mice^56^ were download from SRA (fasterq-dump, sratools v3.0.3) and trimmed using fastp (v0.23.3)^106^ to remove adapters and trailing bases with quality <20. Reads were aligned to GRCm39 using bowtie2 (v2.5.1) in local mode and filtered using samtools (v1.16.1) to retain only properly paired alignments with a quality score of ≥30. PCR duplicates were marked using sambamba (v1.0.1).

#### Visualisation

Heatmaps were plotted using deeptools (v3.5.1)^110^. Coverage tracks were created with deeptools bamCoverage (v3.5.1) with bin sizes of 10 bp or 1 bp and CPM normalisation. For ATAC-seq data, fragments of <100 bp were used to produce coverage tracks corresponding to accessible regions. Tracks at example loci were visualised using the IGV genome browser (v2.9.2)^119^.

The nucleosome positioning plots in Figure 5h were produced by calculating coverage of nucleosome-sized fragments (180-250 bp) around CTCF peaks (centred on CTCF motifs) using deeptools computeMatrix. Mean coverage was calculated in R and plotted using ggplot2. Flanking nucleosome positions were calculated by identifying signal maxima in a window covering 100-300 bp distance from the CTCF motif in both directions.

### Oligonucleotides and antibodies

### Data availability

The UCSC genome browser gRNA track BED file is available from https://hgdownload.soe.ucsc.edu/gbdb/mm10/crisprAll/crispr.bb. CTCF ChIP-seq data (Rep1: SRR6512735, SRR6512736; Rep2: SRR6512737, SRR6512738; Input: SRR6512739, SRR6512740) and ATAC-seq data (SRR6512723, SRR6512724, SRR6512725, SRR6512726) were downloaded from the NCBI SRA (SRP131401, corresponding to GEO accession GSE109671). JASPAR 2024 CORE vertebrate non-redundant data are available from https://jaspar.elixir.no/downloads/. GO annotations for use in Perseus were downloaded from http://annotations.perseus-framework.org (mainAnnot.mus_musculus.txt). The GO Ontology database used within PANTHER is identifiable with DOI: 10.5281/zenodo.8436609. Data from UniProt (https://www.uniprot.org/) and the MGI Gene Ontology Browser (https://www.informatics.jax.org/function.shtml) were used for additional manual annotations where indicated. All other relevant data supporting the key findings of this study are available within the article and its Supplementary Information files, or from the corresponding authors upon reasonable request.

## Acknowledgements

We thank the MRC LMS/National Institute for Health Research Imperial Biomedical Research Centre Flow Cytometry Facility; the MRC LMS Microscopy, Genomics, Proteomics and Bioinformatics facilities; and the MRC LMB Mass Spectrometry Facility for support. We thank Shreya Jha, Nehir Nebioglu and George Young for assistance with ATAC-seq data processing and analysis. We thank Dorus Gadella, Ralf Kuehn, Stefan Stricker, Bradley Cobb and Stephen Smale for gifting reagents. This work was funded by a Kay Kendall Leukaemia Fund Junior Research Fellowship (KKL1334) awarded to A.D., and by support provided by the Medical Research Council UK (MC_UP_1605/12, MC_PC_23024, MC_PC_22015 and MC_UP_1605/11 awarded to A.G.F and M.M. respectively). D.D. was supported by a Wellcome Trust Institutional Strategic Support Fund Springboard award (WCMA_PSN102). For the purpose of open access, the authors have applied a CC BY public copyright licence to any Author Accepted Manuscript version arising from this submission.

## Author contributions

A.G.F. and A.D. conceived the study, wrote the manuscript, and supervised the work. A.D. conducted most of the experiments, analysed data and produced the figures. D.H.G. engineered the PBK KO cell lines and performed additional experiments. E.I-S analysed the ATAC-seq data, C.W. assisted with microscopy and performed image analysis, B.P. isolated mitotic chromosomes by flow cytometry, and S.C. and K.B. performed additional experiments. H.K. and P.S. performed LC-MS/MS proteomics on chromosome and IP samples respectively, and A.M. analysed these data. D.D. contributed to study design, assisted with experiments, and helped to identify ADNP as a bookmarking factor. M.M. provided input and guidance for the study and, as well as J.M.V., supported data analysis and the interpretation of results.

