## Supplementary Figures & Legends for "PBK/TOPK mediates Ikaros, Aiolos and CTCF displacement from mitotic chromosomes and alters chromatin accessibility at selected C2H2-zinc finger protein binding sites"

Dimond, A. *et al.*

Supplementary Figures S1-S6

Legends for Supplementary Videos S1-S4

Supplementary Figure S1

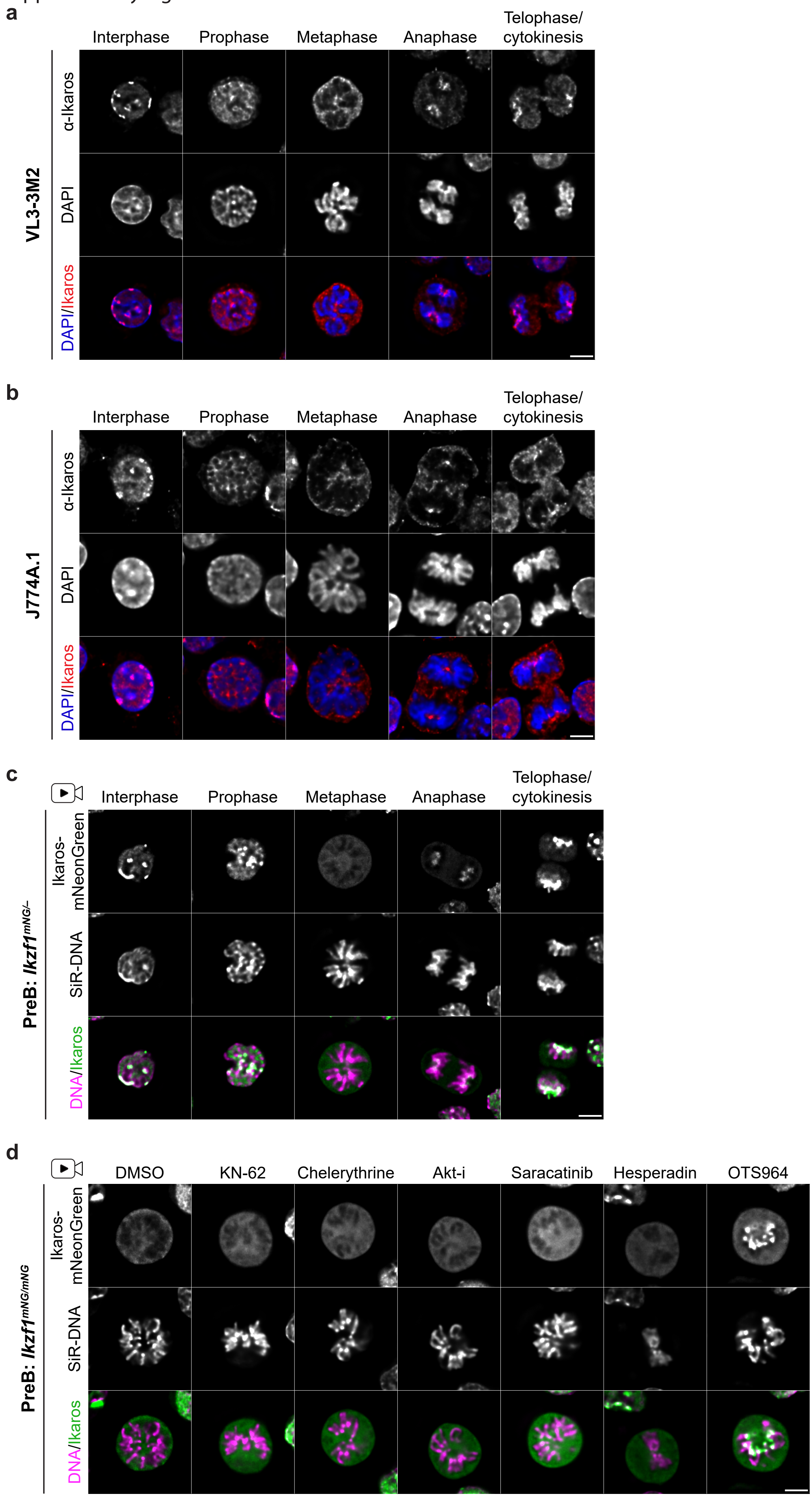

Supplementary Figure S1

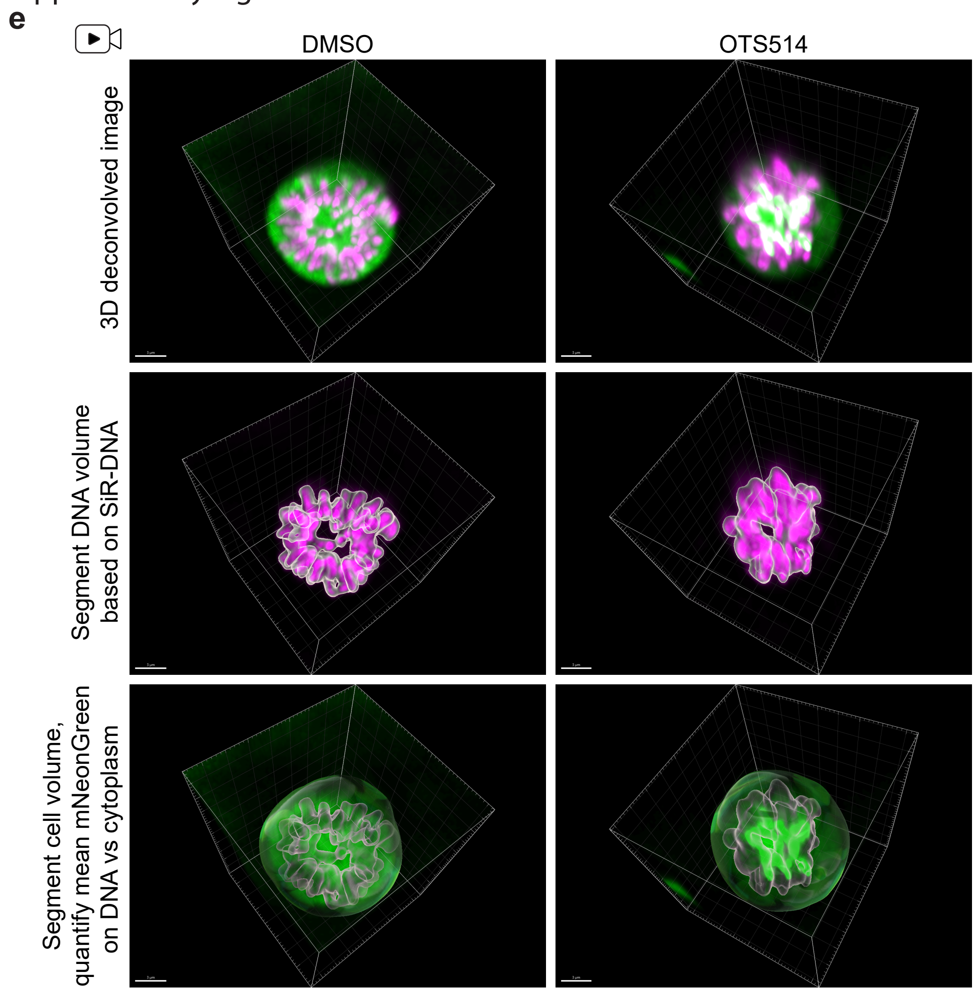


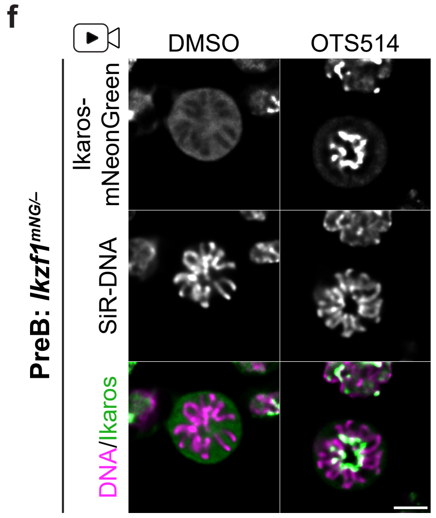

**Supplementary Figure S1 | Ikaros dissociates from metaphase chromosomes in lymphoid and myeloid lineages, but is retained following specific kinase inhibition.**

a. Immunofluorescence staining of Ikaros localisation in interphase and through mitosis in fixed VL3-3M2 cells (mouse T cell line). Images are representative of at least seven cells per mitotic stage (>20 for metaphase) across two independent staining experiments. Scale bar=5 µm.

b. Immunofluorescence staining of Ikaros localisation in interphase and through mitosis in fixed J774A.1 cells (mouse macrophage cell line). Images are representative of at least five cells per mitotic stage (>15 for metaphase) across two independent staining experiments. Scale bar=5 µm.

c. Live-cell images of Ikaros-mNeonGreen in interphase and through mitosis in *Ikzf1*$^{mNeonGreen/–}$ heterozygous KI mouse preB cells (clone 1.2 from Figure 1b) cultured with SiR-DNA. Images are representative of at least six cells per mitotic stage (>20 for metaphase), collected across three independent imaging experiments. Scale bar=5 µm.

d. Live-cell images of *Ikzf1*$^{mNG/mNG}$ mitotic KI mouse preB cells (clone 2.1) showing Ikaros-mNeonGreen localisation following 10 min treatment with 10 µM of the indicated inhibitors. Cells were pre-cultured with SiR-DNA; scale bar=5 µm. Images are representative of at least two independent treatment experiments, with a minimum of 12 mitotic cells imaged per treatment in total.

e. Strategy for quantifying mean chromosomal and cytoplasmic Ikaros-mNeonGreen signal, shown for representative DMSO and OTS514-treated *Ikzf1*$^{mNG/mNG}$ mitotic mouse preB cells. Cropped and deconvolved z-stack images were opened in Imaris software (upper). SiR-DNA signal was used to segment chromosomes (middle) and total cell volume was segmented by background mNeonGreen signal (bottom). Segmented volumes were used to construct a cell in Imaris software, allowing chromosomal and cytoplasmic mNeonGreen mean intensities to be measured separately and expressed as a ratio. Scale bars=3 µm.

f. Representative live-cell images of mitotic *Ikzf1*$^{mNG/–}$ mouse preB cells (clone 1.2) following 10 min treatment with DMSO or 10 µM OTS514. Cells were pre-cultured with SiR-DNA; scale bar=5 µm. Images are representative of >25 mitotic cells from across two independent treatment experiments.

Supplementary Figure S2

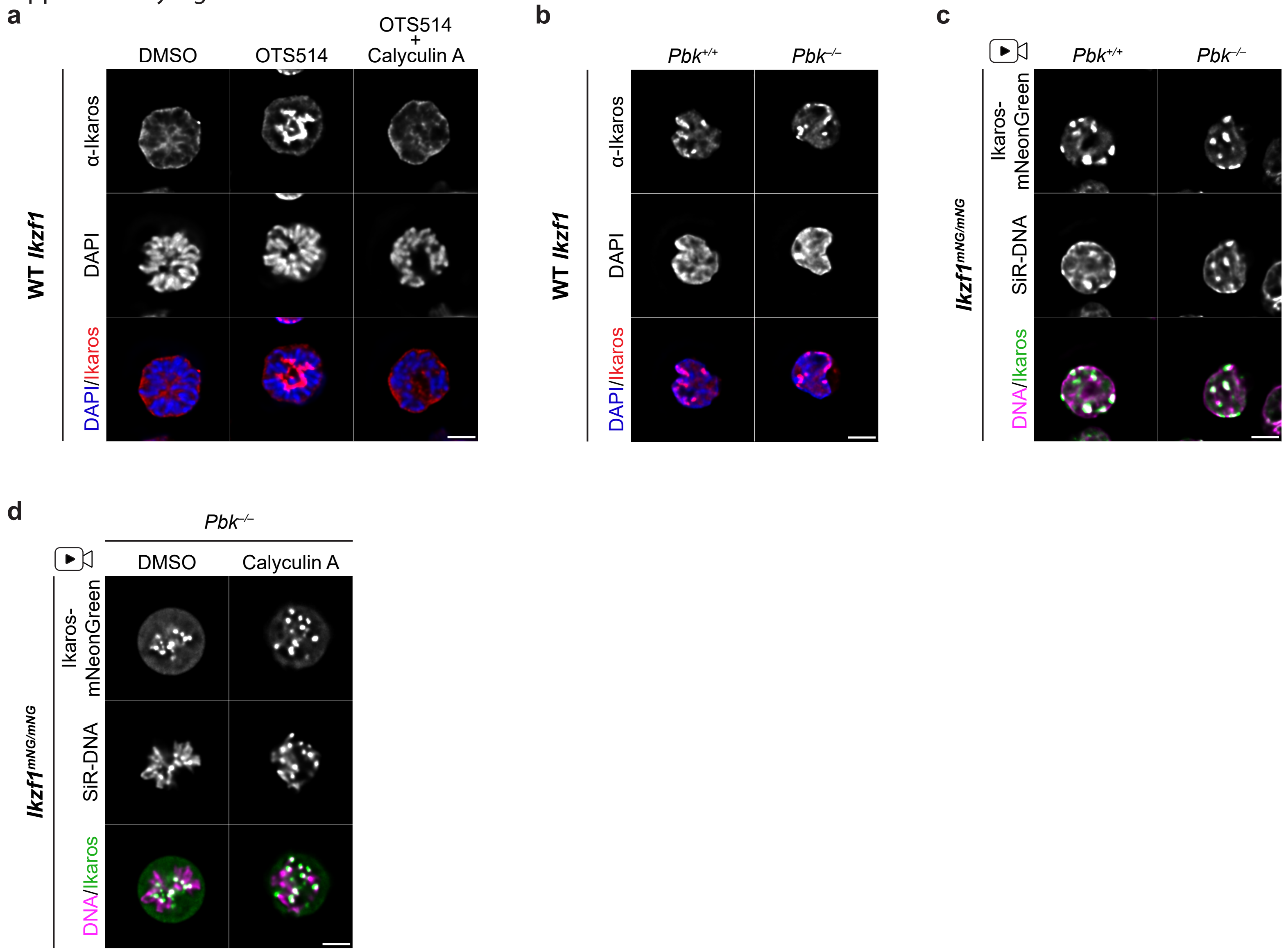

**Supplementary Figure S2 | PBK activity is required for Ikaros dissociation in mitosis.**

a. Immunofluorescence staining of untagged Ikaros localisation in fixed mitotic preB cells following 10 min treatment with DMSO or with 10 µM OTS514 alone or in combination with 100 nM Calyculin A. Images are representative of >18 treated mitotic cells from two independent experiments. Scale bar=5 µm.

b. Immunofluorescence staining of untagged Ikaros showing association with heterochromatin foci in both *Pbk*$^{+/+}$ and *Pbk*$^{-/-}$ interphase mouse preB cells. Images are representative of interphase cells from three independent staining experiments. Scale bar=5 µm.

c. Live-cell imaging of *Pbk*$^{+/+}$ and *Pbk*$^{-/-}$ interphase mouse *Ikzf1*$^{mNG/mNG}$ preB cells cultured with SiR-DNA. Images are representative of interphase cells from three independent imaging experiments. Scale bar=5 µm.

d. Live-cell imaging of *Ikzf1*$^{mNG/mNG}$ *Pbk*$^{-/-}$ mitotic mouse preB cells treated for 10 min with DMSO or 100 nM Calyculin A. Cells were pre-cultured with SiR-DNA; scale bar=5 µm. Images are representative of four independent replicates. All Calyculin A treated mitotic cells imaged (36/36) showed clear foci of mNeonGreen signal at centromeres, despite visible loss of chromosome spindle attachment in many cells.

Supplementary Figure S3

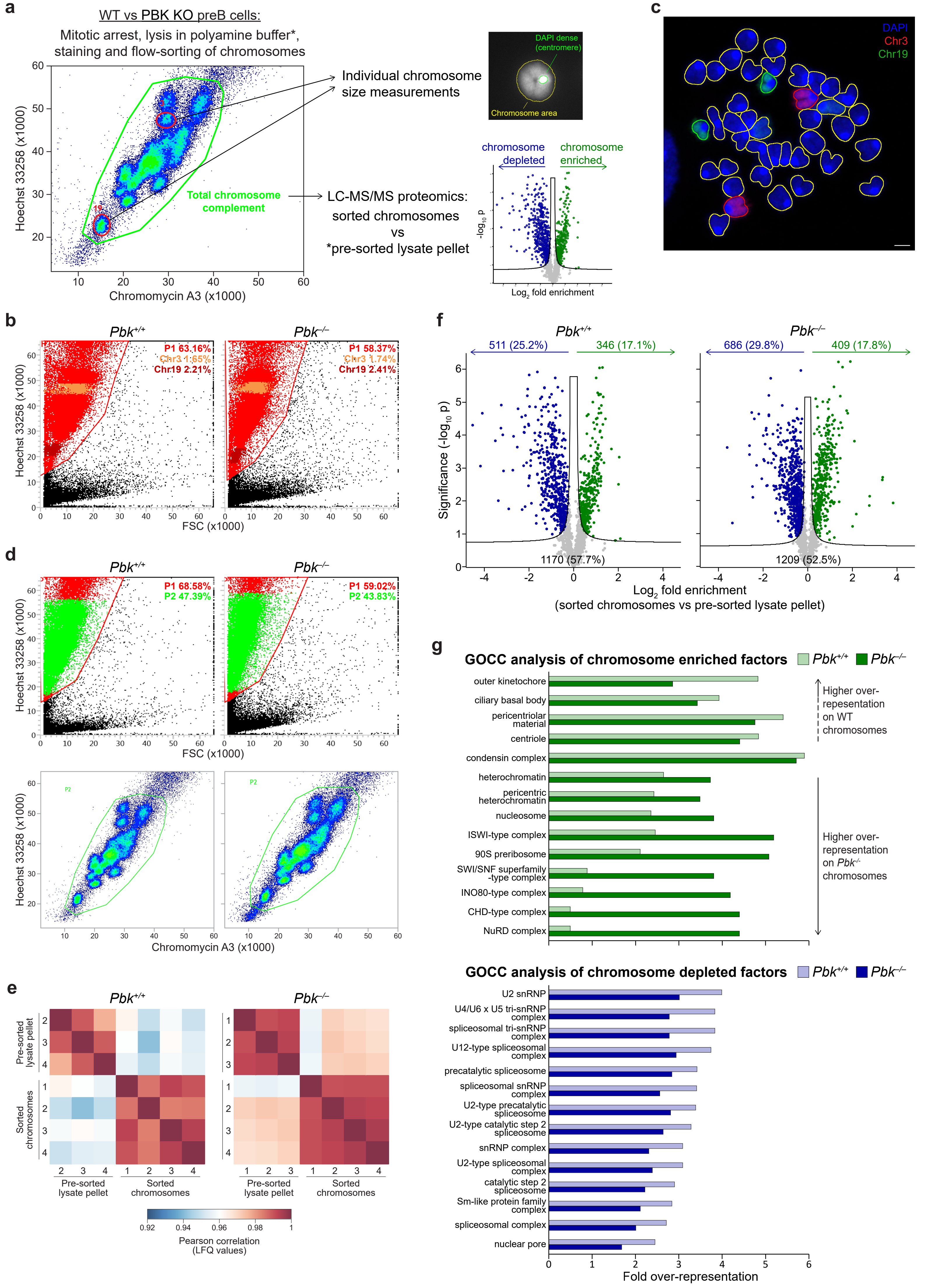

**Supplementary Figure S3 | Analysis of mitotic chromosome size and protein composition in *Pbk*$^{+/+}$ and *Pbk*$^{-/-}$ mouse preB cells.**

a. Strategy for purifying individual or total native mitotic chromosomes by flow-cytometry for size measurements or LC-MS/MS proteomics analysis respectively. Mitotically arrested cells were lysed, chromosomes were released into polyamine buffer and stained with Hoechst 33258 and Chromomycin A3 allowing chromosomes to be visualised and purified by flow cytometry. For size measurements, individual chromosomes 3 and 19 were isolated, cytospun onto poly-L-lysine slides and stained with DAPI, allowing total and DAPI-dense (centromeric) areas to be measured. For proteomics, total purified chromosomes were analysed by LC-MS/MS and compared to unpurified pellets from the lysis step to calculate chromosomal enrichment or depletion of factors.

b. P1 gating strategy for isolation of individual chromosomes 3 and 19. Gates and percentages of total events are shown for a representative experiment, corresponding to Figure 3a.

c. Representative metaphase spread from mouse preB cells (example is from *Pbk*$^{-/-}$ cells) hybridised with chromosome painting probes for chromosomes 3 (red) and 19 (green). Chromosome outlines were segmented on DAPI signal (blue) and filtered based on chromosome paint signals. Scale bar=2 µm.

d. Gating strategy for flow-sorting total chromosomes. Gates and percentages of total events are shown for a representative experiment.

e. Pearson's correlations of proteomics samples after LFQ normalization, showing clear segregation of pre-sorted and sorted samples from both of *Pbk*$^{+/+}$ and *Pbk*$^{-/-}$ cells (rather than by replicate). Note that *Pbk*$^{+/+}$ pre-sorted lysate pellet replicate 1 and *Pbk*$^{-/-}$ pre-sorted lysate pellet replicate 4 were excluded for technical reasons (see Methods).

f. Volcano plots of factors enriched (green) or depleted (blue) from *Pbk*$^{+/+}$ (left) and *Pbk*$^{-/-}$ (right) mitotic chromosomes, compared to pre-sorted lysate pellets (modified two-tailed t-test with permutation-based false discovery rate (FDR)<0.05 and S0=0.1; n=4 chromosome samples and n=3 lysate pellet samples).

g. GO term (cellular component) overrepresentation analysis amongst factors enriched (upper, green) or depleted (lower, blue) on mitotic chromosomes. Analysis was performed separately for *Pbk*$^{+/+}$ (pale green/blue) or *Pbk*$^{-/-}$ (solid green/blue) data, using the total proteins detected in each condition as background. Displayed are the top most overrepresented terms with FDR<0.05, in either condition, ordered by the difference in fold overrepresentation between conditions.

Supplementary Figure S4

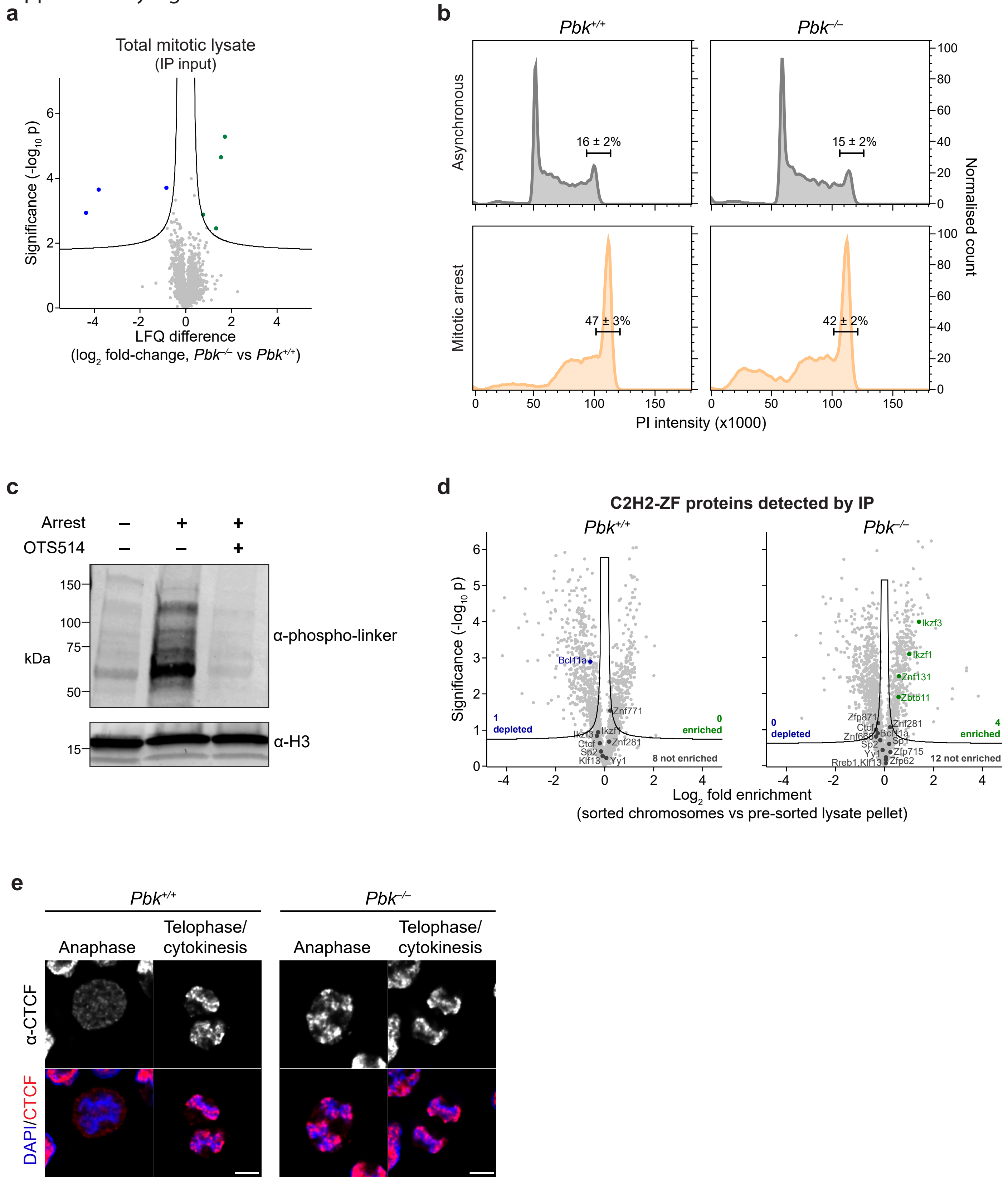

**Supplementary Figure S4 | Loss of PBK activity alters mitotic phosphorylation status and localisation of C2H2-ZF proteins rather than abundance.**

a. Volcano plot comparing protein abundances (LC-MS/MS) in total mitotic lysates from *Pbk*$^{+/+}$ and *Pbk*$^{-/-}$ mouse preB cells, used as inputs for IP in Figure 4c. Seven factors showed significantly different abundances (green/blue; modified two-tailed t-test with permutation-based false discovery rate (FDR)<0.05, S0=0.1; n=3), two further factors (including PBK) were only detected in *Pbk*$^{+/+}$ lysates and eight factors were only detected in *Pbk*$^{-/-}$ lysates; however, none were C2H2-ZF proteins.

b. Representative propidium iodide (PI) profiles of asynchronous (upper, grey) and mitotically arrested (lower, orange) *Pbk*$^{+/+}$ (left) and *Pbk*$^{-/-}$ (right) mouse preB cells. Average percentage of cells (± standard deviation) in G2/M are given for three replicates.

c. Western blot showing loss of phospho-linker detection in mitotically arrested WT mouse preB cells following 10 min treatment with 10 µM OTS514. Total H3 was used as a loading control; representative of two independent replicates.

d. Volcano plots of factors enriched on or depleted from *Pbk*$^{+/+}$ and *Pbk*$^{-/-}$ mitotic chromosomes, as in Figure 3e and Supplementary Figure S3f, highlighting C2H2-ZF proteins identified in Figure 4c as likely PBK targets (green=enriched, blue=depleted, dark grey=not significantly enriched/depleted; modified two-tailed t-test with permutation-based false discovery rate (FDR)<0.05, S0=0.1; n=4 chromosome samples and n=3 lysate pellet samples).

e. Immunofluorescence staining of CTCF in fixed WT (*Pbk*$^{+/+}$) or *Pbk*$^{-/-}$ preB cells in anaphase and telophase/cytokinesis. Representative of cells from two independent staining experiments. Scale bar=5 µm.

Supplementary Figure S5

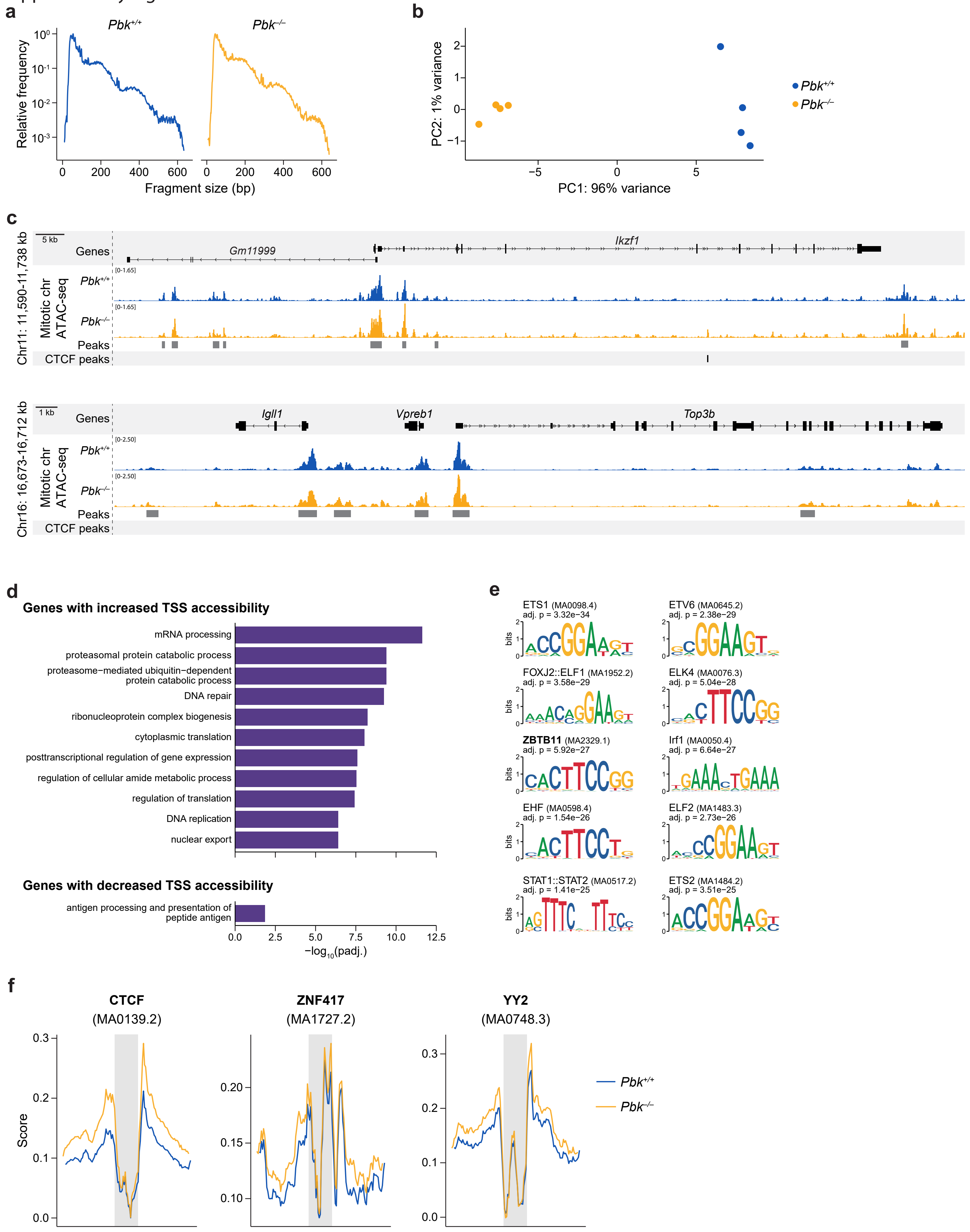

**Supplementary Figure S5 | ATAC-seq analysis of purified mitotic chromosomes from *Pbk*$^{+/+}$ and *Pbk*$^{-/-}$ mouse preB cells.**

a. Fragment length distribution in merged *Pbk*$^{+/+}$ and *Pbk*$^{-/-}$ mitotic chromosome ATAC-seq libraries after genome alignment (n=4+4, plotted on a log10 scale, smoothed by taking the rolling average over 5 bp windows).

b. Principal component analysis based on variance-stabilising transformed read counts in consensus MACS2 peaks.

c. Representative loci showing nucleosome-free (<100bp) ATAC-seq signal from *Pbk*$^{+/+}$ (blue) and *Pbk*$^{-/-}$ (orange) mitotic chromosomes (normalised merged signal for n=4+4). For each locus, gene annotations are shown above; MACS2 consensus ATAC-seq peaks are shown below (none significantly altered in these regions), along with any CTCF peaks from published asynchronous data[56].

d. GO term enrichment (biological process) amongst genes with increased (upper; top ten most significant terms only) or decreased (bottom) TSS accessibility.

e. Top ten enriched motifs in peaks which show decreased accessibility in *Pbk*$^{-/-}$ mitotic chromosomes. ZBTB11, indicated in bold, is the sole C2H2-ZF protein associated with these motifs.

f. ATAC-seq footprint profiles for three of the motifs showing the largest changes in footprint score between *Pbk*$^{+/+}$ and *Pbk*$^{-/-}$ mitotic chromosomes (one representative CTCF motif is shown). Smoothed TOBIAS score (representing cut site density, corrected for Tn5 sequence bias) is plotted ±60 bp from the centre of the indicated motif (grey shading), smoothed by taking the rolling average over 5 bp windows.

Supplementary Figure S6

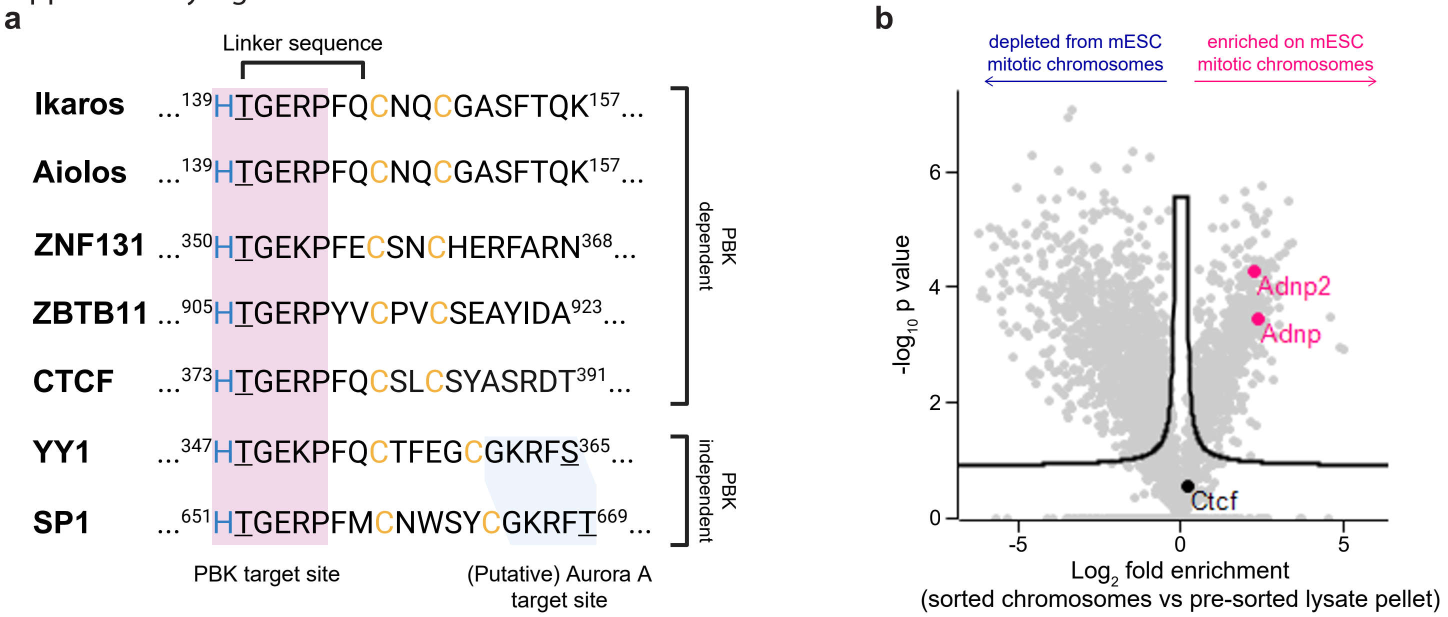

**Supplementary Figure S6 | Comparisons of selected C2H2-ZF proteins.**

a. Sequence comparisons of selected C2H2-ZF proteins which are dependent on or are independent of PBK for mitotic dissociation. Known or putative phosphorylation target sites for PBK and Aurora kinase A are highlighted. YY1 is phosphorylated at the highlighted ZF serine 365 by Aurora kinase A[18]. A closely matching motif is also present in SP1, similarly positioned within a ZF, but is absent from the other factors (either within the ZFs or at other locations).

b. LC-MS/MS proteomic analysis of mESC mitotic chromosomes taken from[8], highlighting CTCF (not significantly enriched) and ADNP/ADNP2 (significantly enriched); modified two-tailed t-test with permutation-based FDR<0.01, S0=0.1, n=3.

**Supplementary Videos S1-4 | Live-cell imaging of Ikaros-mNeonGreen through mitosis.**

*Corresponding to Figures 1b-c and Supplementary Figure S1c.*

Time-lapse live-cell imaging of Ikaros-mNeonGreen (middle panel, green) localisation through mitosis at 3 min intervals in KI mouse preB cell clones pre-incubated with SiR-DNA (left panel, magenta). Video S1=clone 1.1; Video S2=clone 1.2; Video S3=clone 2.1; Video S4=clone 2.2; scale bars=5 µm.
